# Three-dimensional genome architecture persists in a 52,000-year-old woolly mammoth skin sample

**DOI:** 10.1101/2023.06.30.547175

**Authors:** Marcela Sandoval-Velasco, Olga Dudchenko, Juan Antonio Rodríguez, Cynthia Pérez Estrada, Marianne Dehasque, Claudia Fontsere, Sarah S.T. Mak, Valerii Plotnikov, Ruqayya Khan, David Weisz, Vinícius G. Contessoto, Antonio B. Oliveira Junior, Achyuth Kalluchi, Arina D. Omer, Sanjit S. Batra, Muhammad S. Shamim, Neva C. Durand, Brendan O’Connell, Alfred L. Roca, Andreas Gnirke, Isabel Garcia-Treviño, Rob Coke, Joseph P. Flanagan, Kelcie Pletch, Aurora Ruiz-Herrera, Eric S. Lander, M. Jordan Rowley, José N. Onuchic, Love Dalén, Marc A. Marti-Renom, M. Thomas P. Gilbert, Erez Lieberman Aiden

## Abstract

Ancient DNA (aDNA) sequencing analysis typically involves alignment to a modern reference genome assembly from a related species. Since aDNA molecules are fragmentary, these alignments yield information about small-scale differences, but provide no information about larger features such as the chromosome structure of ancient species. We report the genome assembly of a female Late Pleistocene woolly mammoth (*Mammuthus primigenius*) with twenty-eight chromosome-length scaffolds, generated using mammoth skin preserved in permafrost for roughly 52,000 years. We began by creating a modified Hi-C protocol, dubbed PaleoHi-C, optimized for ancient samples, and using it to map chromatin contacts in a woolly mammoth. Next, we developed “reference-assisted 3D genome assembly,” which begins with a reference genome assembly from a related species, and uses Hi-C and DNA-Seq data from a target species to split, order, orient, and correct sequences on the basis of their 3D proximity, yielding accurate chromosome-length scaffolds for the target species. By means of this reference-assisted 3D genome assembly, PaleoHi-C data reveals the 3D architecture of a woolly mammoth genome, including chromosome territories, compartments, domains, and loops. The active (A) and inactive (B) genome compartments in mammoth skin more closely resemble those observed in Asian elephant skin than the compartmentalization patterns seen in other Asian elephant tissues. Differences in compartmentalization between these skin samples reveal sequences whose transcription was potentially altered in mammoth. We observe a tetradic structure for the inactive X chromosome in mammoth, distinct from the bipartite architecture seen in human and mouse. Generating chromosome-length genome assemblies for two other elephantids (Asian and African elephant), we find that the overall karyotype, and this tetradic Xi structure, are conserved throughout the clade. These results illustrate that cell-type specific epigenetic information can be preserved in ancient samples, in the form of DNA geometry, and that it may be feasible to perform de novo genome assembly of some extinct species.

## Introduction

The field of ancient DNA studies began with the sequencing of short organelle fragments from historic samples (Higuchi et al. 1984; Pääbo 1985), and has undergone a remarkable expansion in the subsequent forty years. Current paleogenomic analyses involve DNA sequencing of whole genomes (aDNA-Seq) from a plethora of extinct (e.g. Miller et al. 2008; Green et al. 2010) and extant humans, animals (e.g. Orlando et al. 2013; Rasmussen et al. 2010), plants (e.g. Ramos-Madrigal et al. 2016), and pathogens (e.g. Bos et al. 2011; Martin et al. 2013; Smith et al. 2014), spanning back over one million years (van der Valk et al. 2021). Yet the fragmentary nature of aDNA molecules has restricted such analyses to those based on the mapping of short reads to a modern reference genome. Typically this includes the identification of SNPs and small indels, for use in phylogenomic and population genomic analyses, as well as the identification of variants with likely functional consequences (e.g. Miller et al. 2008; Green et al. 2010; Rasmussen et al. 2010).

Although much has been learned from these methods, the short length of typical aDNA molecules has limited the information that can be gleaned from ancient samples. They overlook larger scale differences, such as chromosomal rearrangements. In fact, the only large scale differences that can be identified are cases where a sequence from the modern reference dataset is absent in the ancient genome (Martin et al. 2013). While for modern samples, larger scale sequence features can be explored using techniques based on the recovery of high molecular weight (HMW) DNA, this dependency on HMW DNA is an obstacle when using ancient samples.

Ancient DNA studies have also explored epigenetic differences, such as by recovering methylated cytosines from ancient templates (Llamas et al. 2012; Pedersen et al. 2014; Gokhman et al. 2014) or through differential decay signatures, which are proposed to relate to DNA placed within and outside of nucleosomes (Pedersen et al. 2014). Yet there is no genome-wide epigenetic data for many ancient species of interest, such as the woolly mammoth. Here too the short length typical of aDNA makes it more difficult to reliably place epigenetic features in their correct genomic context.

One class of techniques that have proven useful for generating larger-scale insights into genomes are those based on Hi-C (Lieberman-Aiden et al. 2009), which interrogates the shape of whole chromosomes. Hi-C uses DNA-DNA proximity ligation (Cullen, Kladde, and Seyfred 1993; Dekker et al. 2002) to create chimeric reads derived from DNA strands that are close to one another in the 3D space within the cell nucleus, even if these DNA strands are distant along the contour of the chromosome.

Contact frequencies derived from Hi-C have many uses. For instance, they can be used to determine the relative positioning of sequences along chromosomes, insofar as sequences that tend to be in contact are more likely to be near one another in the 1D genome sequence. Consequently, Hi-C datasets can be used to facilitate genome assembly, to find errors in contigs (contiguous sequences) that are incorrectly assembled, and to reliably order and orient contigs into chromosome-length scaffolds (Burton et al. 2013; Kaplan and Dekker 2013; Dudchenko et al. 2017). Hi-C maps have also been shown to reflect the epigenetic state and activity levels of loci genome-wide (Lieberman-Aiden et al. 2009; Rao et al. 2014). More broadly, Hi-C maps provide a wealth of information about the architecture of the entire genome in 3D. Generation of Hi-C datasets for ancient samples would therefore be quite valuable.

Although to date Hi-C has been applied in settings where there is HMW DNA in the sample being processed, we hypothesized that HMW DNA may not be an absolute requirement, and that degraded DNA could still yield a suitable substrate for Hi-C so long as the overall 3D conformation of chromosomes is maintained (Figure 1A). To test this hypothesis we developed PaleoHi-C, a variant of the in situ Hi-C protocol (Rao et al. 2014) specifically adapted for ancient samples (Figure 1B). We demonstrated PaleoHi-C by applying it to a permafrost-preserved woolly mammoth skin sample, thereby generating a catalog of DNA-DNA contacts in woolly mammoth skin nuclei. These results highlight new approaches to the study of genomic architecture, epigenetics, and transcriptional state in ancient samples, and open the door to generating de novo genome assemblies for ancient and extinct species.

**Figure 1.**
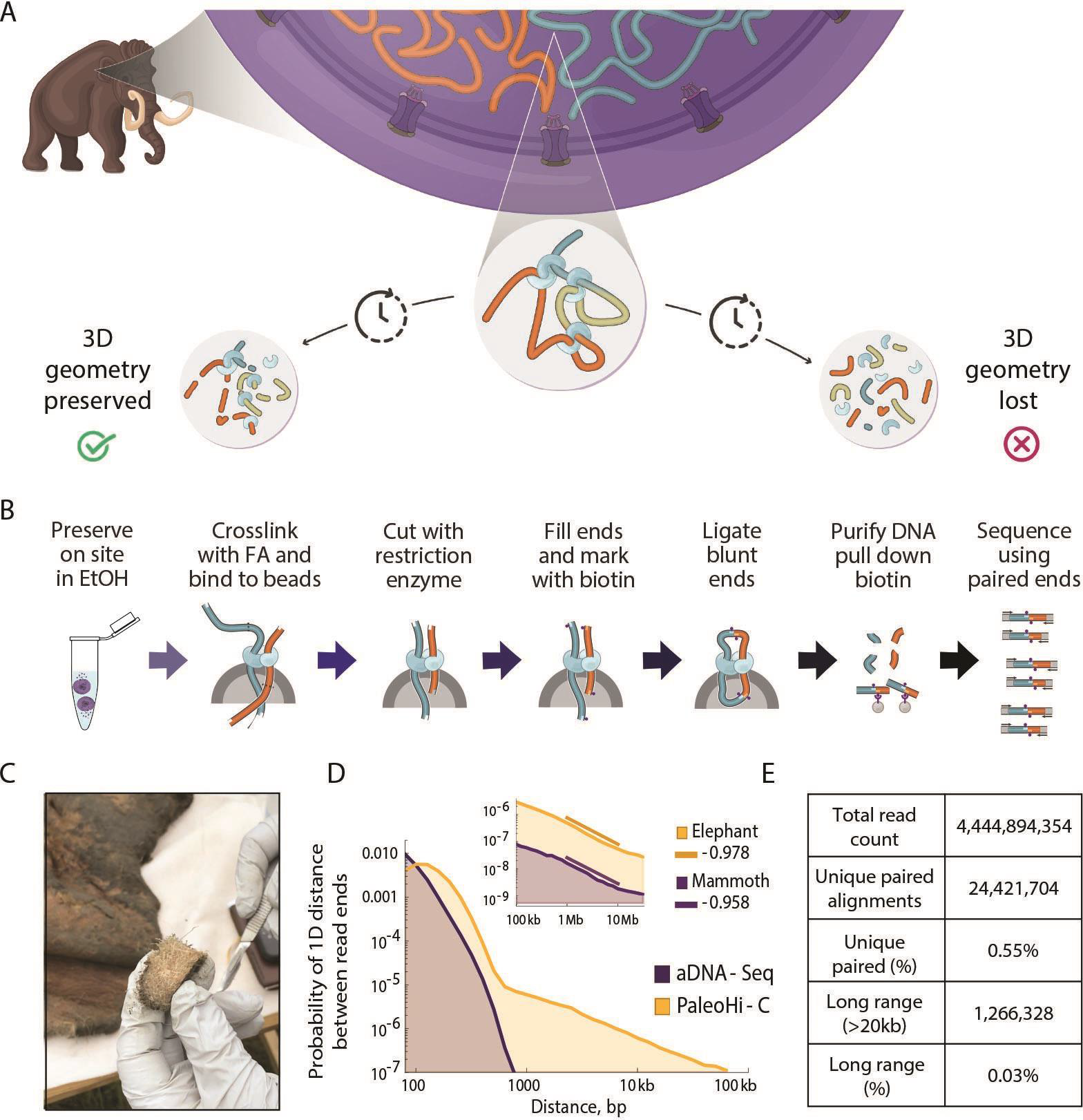
PaleoHi-C reveals that the geometry of DNA is preserved in a 52,000-year-old sample of woolly mammoth skin. **A**. DNA degradation is a common occurrence in paleo-samples, which can pose significant challenges for researchers seeking to extract genetic information from ancient remains. As long as the overall 3D conformation of the chromosomes is maintained, degraded DNA could still yield a suitable substrate for Hi-C: DNA-DNA proximity ligation coupled with sequencing. **B**. Overview of PaleoHi-C. Samples were collected into ethanol in the field. The laboratory procedure begins when the samples are crosslinked with formaldehyde (FA) and bound to beads, where they are cut with a restriction enzyme. The ends are repaired and blunted while introducing biotin. DNA-DNA-proximity ligation is performed, and the ligation junctions are captured on streptavidin beads and sequenced. All steps are performed in small volumes on account of the small amounts of genomic material. **C**. A photograph of the mammoth sample used in this study. **D**. PaleoHi-C retrieves long-range interactions preserved in the Late Pleistocene sample. Histograms on a log-log plot showing the 1D genomic distance between opposite ends of a sequenced read for ordinary aDNA-Seq, and for PaleoHi-C. For PaleoHi-C, a small fraction of the reads reflects Hi-C contacts between loci that lie far away in 1D. No such population is seen in an ordinary aDNA-Seq assay on the same sample. **D, inset**. 1D genomic distance between opposite ends of a sequenced read for PaleoHi-C on woolly mammoth skin vs. in situ Hi-C on Asian elephant skin. For Hi-C experiments, like those shown here, the y-axis corresponds to relative contact probability. A power law is seen in the 1Mb to 10Mb distance range, with a nearly identical scaling (Mammoth: −0.96; Elephant: −0.98), in both experiments. **E**. PaleoHi-C summary statistics.

## Results

### We developed PaleoHi-C and applied it on a 52,000-year-old sample from the skin of a woolly mammoth

We collected a sample from a Late Pleistocene woolly mammoth (†*Mammuthus primigenius*), ID IN18-032, near Belaya Gora, Sakha Republic (N68.57887, E147.16055, Figure 1C). Conventional paleogenomic sequencing data has previously been published from this specimen under the ID “YakInf” (Díez-del-Molino et al. 2023). We dated the sample using both carbon dating, which implied that the sample was at least 45,000 years old (Díez-del-Molino et al. 2023), and analysis of mitochondrial DNA, which suggested that the sample was ca. 52,000 years old (Supplementary Online Information). Upon visual inspection, the skin appeared to be unusually well-preserved, with no signs of putrefaction. Following removal of the outer surface, two subsamples were collected from behind the ear using a scalpel (Figure 1C, Figure S1).

We applied a modified in situ Hi-C protocol (Rao et al. 2014), dubbed PaleoHi-C, to generate chromatin contact data for IN18-032. As compared to in situ Hi-C, the principal changes are: (i) immediate preservation in ethanol to enhance sample resilience and prevent microbial activity; and (ii) omission of fragmentation and size selection after biotin pull-down which is unnecessary given the lack of HMW DNA. See Figure 1B and Supplementary Online Information. A total of 26 sequencing experiments were conducted for the IN18-032 sample, generating ∼4.4 billion reads (Figure 1E, Table S1).

We then plotted the 1D genomic distance separating the alignments derived from a given read pair when aligned to the largest scaffold (GL010027.1) of the published assembly for the African elephant, Loxafr3.0 (Palkopoulou et al. 2018). The results indicate an enrichment of pairs of Hi-C reads for all distances above ∼100 bp as compared to conventional shotgun aDNA sequencing (Figure 1D). However, the number of chromatin contacts was much lower than what is typically obtained for modern samples (Lieberman-Aiden et al. 2009; Rao et al. 2014). For instance, only about 1.3 million read pairs aligned to positions on the same scaffold, but more than 20kb apart along the Loxafr3.0 genome assembly (0.03%), compared to 15.71% seen when using a modern African elephant sample (Table S1, S2). This is consistent with the fact that the ancient mammoth tissues have undergone degradation since the time of death.

Despite this, the key characteristics of chromatin contacts in mammoth appear to be similar to those in modern samples. For example, the frequency of chromatin contacts in mammoth decreased monotonically when compared to 1D distance along the contour of the chromosome. A power law relating contact probability and 1D distance was observed for distances between 1Mb and 10Mb, with a slope of −0.96 (Figure 1D inset, Figure S3). This matches the power law seen when in situ Hi-C is performed on modern Asian elephant skin (slope of −0.98) and closely resembles the decline seen for Hi-C data from modern samples more generally (Lieberman-Aiden et al. 2009; Rao et al. 2014). Notably, this finding is consistent with the preservation of the local, globular structure of woolly mammoth chromatin despite 52,000 years in permafrost and indicates conserved synteny between elephant and mammoth.

In the ancient DNA field, surprising claims have sometimes proven, on closer examination, to be the result of sample contamination (Hofreiter et al. 2001). We therefore sought to preclude the possibility that our PaleoHi-C results were due to contamination of woolly mammoth sample IN18-032 by material from modern elephants. First, we performed PaleoHi-C in an ancient DNA laboratory that has never been used to process modern elephant samples. Second, we used PMDTools (Skoglund et al. 2014) to look for patterns of DNA damage, such as cytosine deamination in single-stranded 5’ overhangs, that are typically elevated in ancient DNA samples (Briggs et al. 2007; Brotherton et al. 2007). We found that levels of DNA damage were significantly elevated in all mammoth libraries as compared to modern elephant libraries (Figure S4A, Table S4). The effect was unchanged when we restricted our analysis to reads corresponding to long-range contacts. Notably, the percentage of reads from IN18-032 carrying damage, although higher than in modern samples, were relatively low for an ancient sample, consistent with the sample’s overall high level of preservation. Finally, we assigned individual reads to a putative species-of-origin, whenever possible, based on single-nucleotide polymorphisms (SNPs) that are fixed in woolly mammoth but absent in modern elephants, or vice versa (Díez-del-Molino et al. 2023). We found that reads from mammoth libraries - when they could be assigned - were overwhelmingly assigned to mammoth (99.8% of the time) and almost never assigned to elephant (0.2%), whereas reads from elephant libraries were overwhelmingly assigned to elephant (99.9%) and almost never assigned to mammoth (0.1%) (Table S5). This remained the case even when we restricted our analysis to reads corresponding to long-range contacts (Figure S4B, Table S5). The few cases of discordant assignments could be explained by sequencing errors. We conclude that the signal in PaleoHi-C experiments derives from ancient mammoth DNA, rather than from modern contaminants.

Taken together, these results suggest (i) that DNA geometry can be well preserved in ancient samples, such as our 52,000-year-old sample of the woolly mammoth; and (ii) that PaleoHi-C is able to extract long-distance chromatin contacts from such ancient samples.

### Reference-assisted 3D genome assembly of a woolly mammoth yields chromosome length scaffolds

To illustrate the uses of PaleoHi-C data, we developed a method that could be used to create a genome assembly with chromosome-length scaffolds using only aDNA-Seq and small quantities of PaleoHi-C data. The method begins by aligning the Hi-C data for a target species to an “assisting” reference genome of a related species. The purpose of this step is to identify large scale changes that arise over the course of evolution (as well as incongruities that simply reflect errors in the assisting genome assembly). Once these changes are identified, they are used to make corresponding modifications to the assisting genome, such as splitting sequences and changing their ordering and orientation. These changes lead to a modified genome assembly whose chromosome-length scaffolds reflect the large-scale structure of the target genome. Next, aDNA-Seq and Hi-C data are compared to the resulting assembly in order to correct the local sequence, yielding a reference genome for the target species with chromosome-length scaffolds. Notably, this strategy can yield chromosome-length scaffolds even if the assisting genome assembly is highly fragmented (Figure 2A).

**Figure 2.**
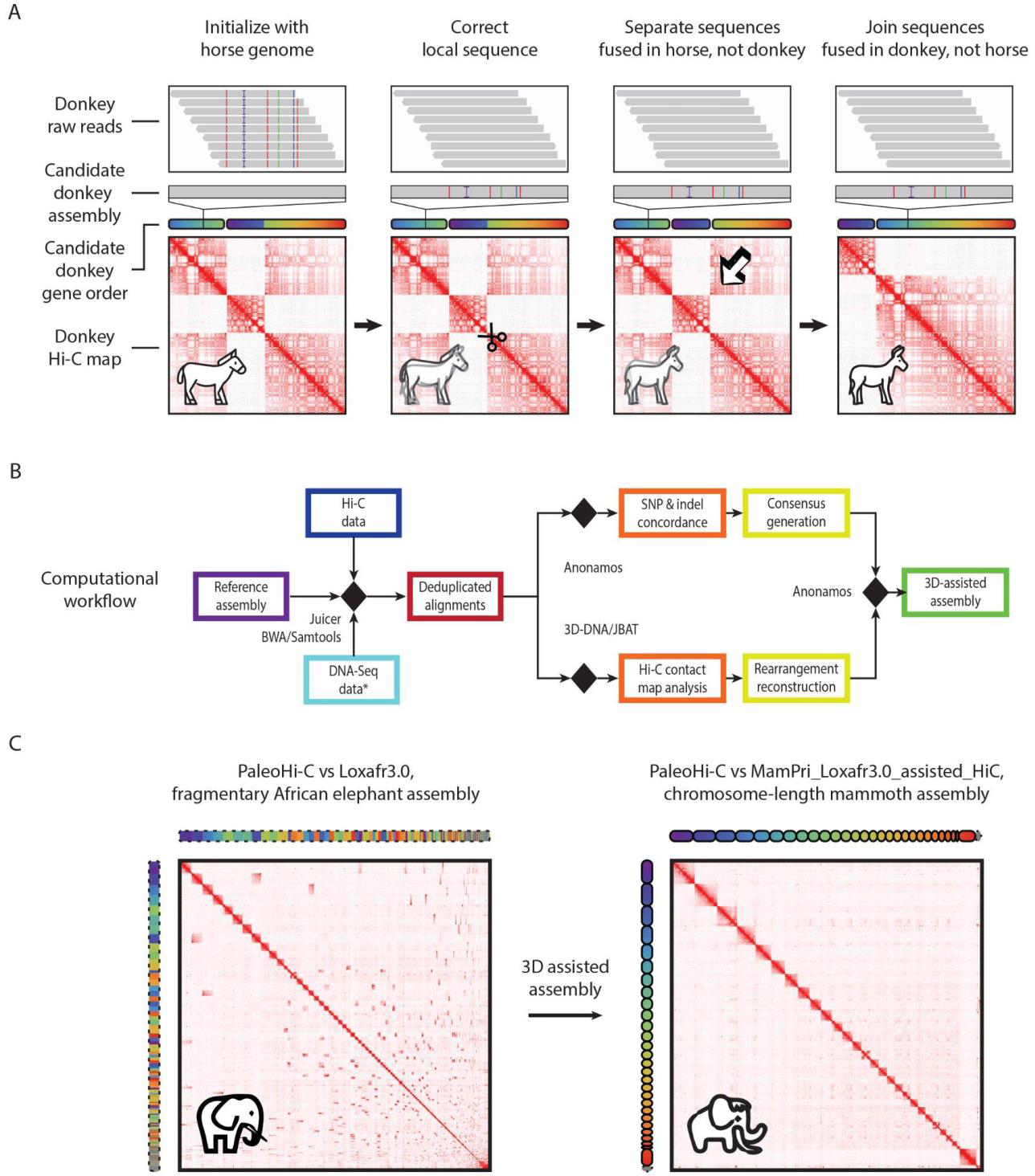
We developed 3D assisted assembly and used it to generate an assembly of the woolly mammoth genome with chromosome-length scaffolds. **A**. Schematic representation of 3D assisted assembly. We illustrate the process by focusing on two sequences in the horse genome, chr8 and chr24, which fuse and break to ultimately become donkey chr9 and a part of donkey chr5. The 3D assisted assembly is shown as a 4-step process, from left to right. We begin by assuming that the donkey genome assembly is identical to the assisting horse genome assembly, EquCab2.0, which we refer to as the “step-1 donkey genome assembly”. The diagrams explaining the step in detail are shown in the leftmost column “Initialize with horse genome”. In successive columns, we modify this initial draft step-by-step, obtaining a genome assembly for the donkey with accurate local sequence and accurate chromosome-length scaffolds. Bottom row: Hi-C contact map showing reads from the donkey aligned to a series of candidate reference genome assemblies for the donkey: step-1, step-2, step-3, and step-4. 2nd to last row: chromogram illustrating the order of loci in a series of candidate reference genome assemblies for the donkey, with respect to the final (step-4, correct) order. Blue is at the beginning of the final order, and red is at the end. Similar colors indicate that the loci are nearby in 1D in the final donkey genome assembly. Changes in the order of loci, and in their partitioning among chromosomes, are shown by means of the chromogram. 3rd row: zoom-in on the DNA sequence at a given locus, for a series of candidate reference genome assemblies for the donkey. Gray coloration indicates that the sequence matches the sequence for the assisting genome assembly, horse EquCab2.0. Colored bars represent single base changes. The ‘I’ icon indicates an insertion. The color indicates the resulting base: A (green), C (blue), G (yellow) and T (red) respectively. Top row: individual donkey sequence reads aligned to the locus depicted in the 3rd row. Differences between the individual reads and the draft are shown using the same color convention as in the 3rd row, except that the colors are now with respect to the draft donkey genome in the same column, rather than with respect to the assisting horse genome. The procedure for 3D assisted assembly is as follows. (Left column). DNA-Seq and Hi-C reads derived from the donkey are aligned to the step-1 donkey genome assembly which is identical to the assisting horse genome assembly, EquCab2.0 (top two rows). The aligned donkey Hi-C reads are used to make a contact map for the donkey with respect to the step-1 donkey genome assembly (bottom row). Local differences between the aligned reads and the step-1 donkey reference are immediately apparent (top two rows). Consequently, individual read alignments are examined and corresponding changes to the step-1 donkey genome assembly are introduced to resolve alignment mismatches. This is similar to the process used in traditional DNA resequencing. The result is the step-2 donkey genome assembly, shown in the 2nd column. Subsequent steps focus on the construction of accurate chromosome-length scaffolds for donkey. This is accomplished by looking for inconsistencies between the donkey Hi-C data and the putative donkey genome assembly. As such, the Hi-C contact map becomes the focus of the assembly process, and is used to iteratively correct the genome assembly scaffolds. (2nd column) In this column, an anomalous feature is highlighted with a pair of scissors: two long sequences which are adjacent to each other on a single chromosome in EquCab2.0 (see chromogram) do not form frequent contacts with one another, indicating that they lie on different chromosomes in the donkey. As such, they should be placed on separate scaffolds, a change that is made in the step-3 donkey genome assembly. (3rd column) The step-3 donkey genome assembly contains three long scaffolds, as shown in the chromogram. Examining the Hi-C map indicates that the first and last of these scaffolds are in frequent 3D contact, implying that they are adjacent in the donkey genome. This change is made in the step-4 donkey genome assembly. (4th column) Both the donkey reads (top) and the Hi-C contact map (bottom) are consistent with the step-4 donkey genome assembly, and the process terminates, with the step-4 genome assembly as the final, 3D assisted genome assembly of the donkey. Of course, in reality, this process is performed on all the donkey chromosomes at once. 3D assisted genome assembly requires extremely small quantities of Hi-C data, making it compatible with PaleoHi-C. It results in a reference that closely matches the true donkey genome both at the single-base level and at the scale of whole chromosomes. **B**. Detailed computational workflow for assisted 3D assembly. DNA-Seq data and/or Hi-C data are aligned to the assisting reference assembly, yielding deduplicated alignments. These alignments are then analyzed with 3D-DNA (Dudchenko et al. 2017) and Juicebox Assembly Tools (Dudchenko et al. 2018) to correct scaffolds and with Anonamos (see Supplementary Online Information) to correct local sequences and incorporate the scaffolding information. Note that DNA-Seq data need not be used if the Hi-C data is sufficiently deep. In this case the reference is corrected, at base-pair resolution, using Hi-C reads alone. **C**. 3D assisted assembly of the woolly mammoth, where we assist using the African elephant draft genome assembly Loxafr3.0 (left). The resulting assembly contains 28 chromosome-length scaffolds. The Hi-C maps for this panel can be browsed interactively at https://tinyurl.com/2ah6rfdz.

We validated our approach by using EquCab2.0 (Wade et al. 2009), a genome assembly for the horse, *Equus caballus*, to assist in the creation of a chromosome-length genome assembly of the donkey, *Equus asinus*. The resulting genome assembly, EquAsi_EquCab2.0_assisted_HiC, comprises 2.25Gb of sequence partitioned among 31 chromosome-length scaffolds (Figure S5, Table S9). We confirmed the accuracy of this strategy by comparing EquAsi_EquCab2.0_assisted_HiC to ASM303372v1_HiC, an assembly of the donkey that we created using the published de novo contig set for the donkey, ASM303372v1 (Renaud et al. 2018), see Figure S6, Table S9. For example, base calls in the EquAsi_assisted_HiC mitochondrial sequence match the de novo assembly ASM303372v1_HiC in 99.6% of positions. Strikingly, this strategy was successful despite the fact that there are numerous large scale rearrangements in the genome of the donkey when it is compared to the genome of the horse (Figure S7). Taken together, these findings confirm that the 3D assisted assembly strategy yields accurate sequences as well as accurate chromosome-length scaffolds.

We then applied this strategy to data for the woolly mammoth sample IN18-032, in combination with shotgun aDNA sequencing data for the same sample (Díez-del-Molino et al. 2023), to produce a genome assembly, MamPri_Loxafr3.0_assisted_HiC, for the woolly mammoth. We assisted using the RefSeq-annotated Loxafr3.0 African elephant genome assembly (Palkopoulou et al. 2018). Despite the fact that Loxafr3.0 is fragmented, the resulting woolly mammoth assembly exhibited 28 chromosome-length scaffolds comprising 3.2 Gb of sequence (98.4% of total sequence), with a contig N50 length of 54kb (Figure 2C and Table S12).

Interestingly, we found that one of the large scaffolds in Loxafr3.0 (GL010038.1, 67,275,361 bp) needed to be split to generate MamPri_Loxafr3.0_assisted_HiC because its sequence was found not be contiguous in the woolly mammoth. To determine whether this reflected a misassembly in Loxafr3.0 or a karyotypic difference between the African elephant and the woolly mammoth, we upgraded fragmented genome assemblies for African elephant (Palkopoulou et al. 2018) and Asian elephant (Tollis et al. 2021) using published in situ Hi-C data (Hoencamp et al. 2021; Álvarez-González et al. 2022) and newly-generated in situ Hi-C data, respectively. We obtained chromosome-length scaffolds in both cases (Figure S8 and S9). In fact, the contig reflected a misassembly in Loxafr3.0. The genome assemblies we obtained for the African elephant, the Asian elephant, and the woolly mammoth showed extremely high conservation of synteny (Figure S10).

To annotate genes in MamPri_Loxafr3.0_assisted_HiC, we ran TOGA (Kirilenko et al. 2023). One way to assess the quality of a gene annotation set is by looking for the fraction of Benchmarking Universal Single Copy Orthologs which are complete; until recently, values in excess of 95% were mostly seen for model organisms (Seppey, Manni, and Zdobnov 2019). We obtained 94.7% BUSCO complete genes (Table S13), demonstrating that reliable gene annotations can be obtained for ancient species using our 3D assisted assembly procedure combined with contemporary gene annotation pipelines such as TOGA.

When aligned to MamPri_Loxafr3.0_assisted_HiC, the PaleoHi-C data - which reflects a total of 1.8 million long-range contacts (>20kb) - facilitates the analysis of the genome architecture and epigenetics of woolly mammoth skin.

### The woolly mammoth exhibits a type II nuclear architecture marked by strong chromosome territories

Eukaryotic genomes tend to fold into one of two architectural types (Hoencamp et al. 2021). Type I is a more polarized architecture, exhibiting features such as telomere clustering, centromere clustering, and chromosomal hairpin structures, which are associated with the arrangement of chromatin along a telomere-to-centromere axis. Type II lacks the above features and is instead marked by prominent chromosome territories, when a chromosome occupies a discrete subvolume of the nucleus, excluding other chromosomes. The PaleoHi-C mammoth data clearly indicates a territorial, or Type II, architecture (Figure 2C), which is commonly observed in mammals with typically sized chromosomes (Hoencamp et al. 2021; Álvarez-González et al. 2022). Although the fact that chromosomes in woolly mammoth occupied distinct territories in the nucleus is not surprising, it is notable that this architectural feature persists after 52,000 years in permafrost. This observation highlights the fact that the overall geometry of chromosomes can be well preserved even while the DNA itself is highly fragmented.

### PaleoHi-C maps reflect cell-type specific genome segregation between the active (A) and inactive (B) compartments, making it possible to nominate genes whose activity is different in mammoth skin as compared to Asian elephant skin

In numerous species, contact maps reveal the segregation of active and inactive chromatin loci to form two compartments that collectively span the genome, dubbed the A and B compartments, respectively. This segregation leads to a plaid pattern in the contact matrix, with elevated contact frequencies between loci in the same compartment, and diminished contact frequencies between loci in different compartments. The plaid pattern can be enhanced by normalizing the contact matrix to account for the effects of 1D sequence proximity, and then calculating the 1st- or 2nd- order autocorrelation matrix (Lieberman-Aiden et al. 2009).

Remarkably, we observed this plaid pattern in PaleoHi-C maps from the woolly mammoth, and successfully enhanced the signal by means of the 2nd-order autocorrelation matrix (Supplemental Online Information). These findings are consistent with the presence of A and B compartments in woolly mammoth skin nuclei, and the retention of their underlying geometry despite 52,000 years in permafrost (Figure 3A, S11, S13).

**Figure 3.**
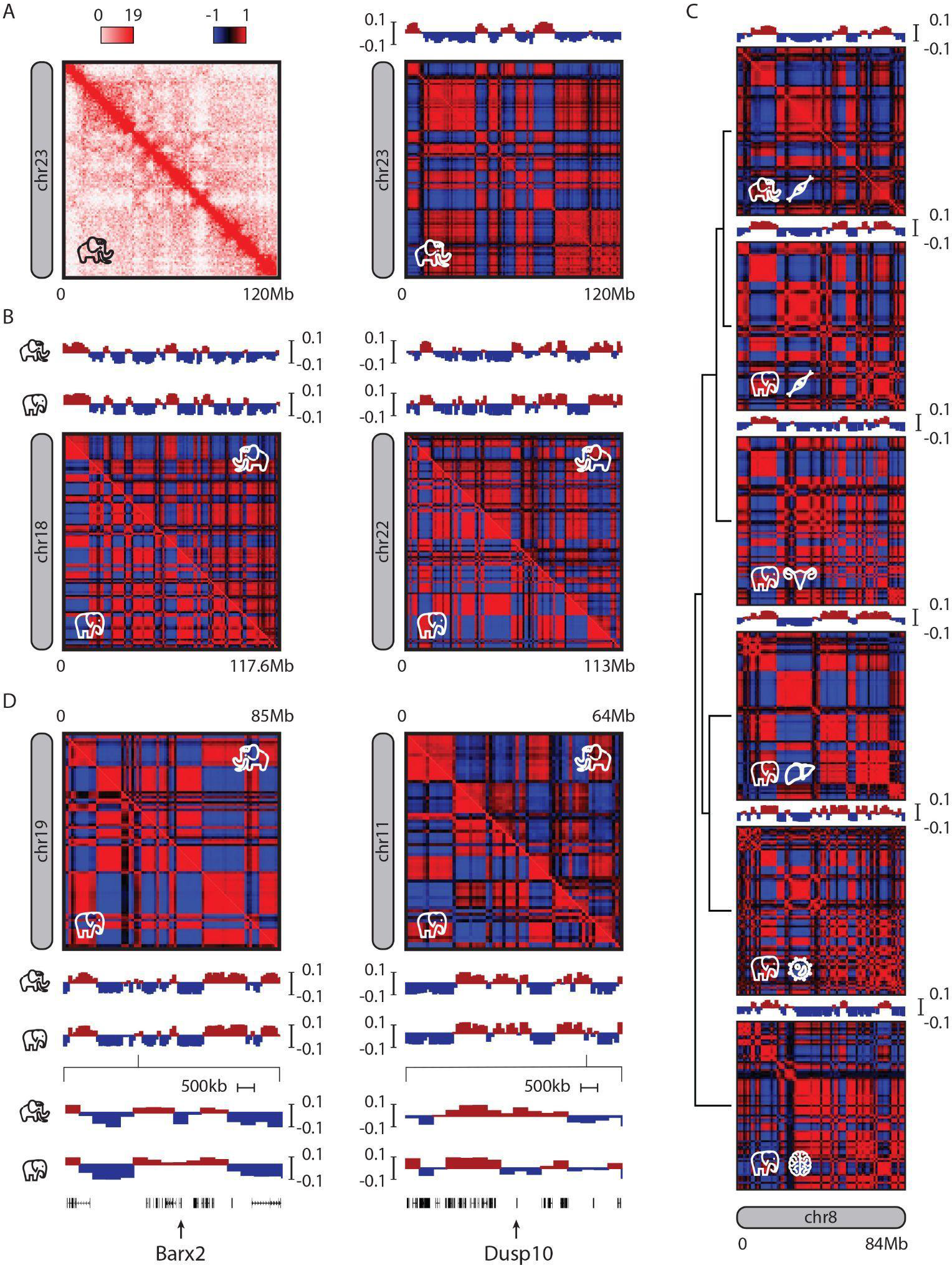
The segregation between the active (A) and inactive (B) genome compartments can survive in ancient samples, making it possible to compare gene activity in woolly mammoth and Asian elephant skin. **A**. PaleoHi-C contact maps from woolly mammoth skin exhibit a plaid pattern, characteristic of the spatial segregation between the A and B genome compartments. For the purpose of comparison to other elephantids, all panels in this and subsequent figures are with respect to the chromosome-length assembly Loxafr3.0_HiC, generated here by using modern Hi-C data (Álvarez-González et al. 2022) to upgrade the published African elephant reference genome, Loxafr3.0 (see Supplementary Online information). Here we show raw contact data corresponding to chromosome 23 in Loxafr3.0_HiC (chromosome 8 in mammoth) as well as the 2nd-order autocorrelation matrix, which enhances the signal and is of great utility when examining low-resolution contact maps. The principal eigenvector of the latter matrix can be used to distinguish between the A (active) and B (inactive) chromatin compartments. Here we adopt the common sign convention of having positive values for the eigenvector corresponding to the A compartment. **B**. The 2nd-order autocorrelation matrices and the corresponding eigenvectors for two representative chromosomes (18 & 22) in both the mammoth and modern Asian elephant skin samples. The maps are highly similar. **C**. PaleoHi-C captures cell-type specific patterns of chromatin activation. Here we show 2nd-order autocorrelation matrices, and the corresponding principal (A/B) eigenvectors, for chromosome 8, for six samples, in the following order from top to bottom: woolly mammoth skin PaleoHi-C; and Asian elephant in situ Hi-C for skin, ovary, liver, blood, and brain. The accompanying dendrogram is based on Euclidean distance between eigenvectors. Woolly mammoth skin is a closer match to modern Asian elephant skin than any of the other modern Asian elephant tissues. **D**. We can use differences in A/B compartment pattern to nominate genes in woolly mammoth skin whose activation state differs from what is seen in Asian elephant skin. Here we show chromosomes 19 and 11. Zoom-in: an apparent difference is seen in the region surrounding genes Barx2 (on chromosome 19) and Dusp10 (on chromosome 11). All maps in this figure are shown at 1Mb resolution.

A and B compartment assignments for individual loci can be determined from a Hi-C map by calculating the first principal component of the contact matrix. We did so for the woolly mammoth contact matrix binned at 1Mb resolution and compared the results to those obtained for a panel of tissues derived from the Asian elephant: skin, ovary, peripheral blood monocytes (PBMCs), liver, and brain. Strikingly, the assignments in elephant skin were consistently more similar to those obtained from mammoth skin than to those obtained from other elephant tissues (Pearson’s r for elephant skin vs. mammoth skin: 0.927; vs. elephant ovary: 0.855; vs. elephant liver: 0.829, vs. elephant brain: 0.804; vs. elephant PBMCs: 0.763, Figure 3B, C). This implies that, despite the extraordinary length of time that has elapsed, the ancient woolly mammoth sample retained cell-type specific architectural features, reflecting patterns of gene activity in the mammoth skin while it was alive.

Reliable compartment assignments can be used to infer the structure of chromosomes in three-dimensional space by means of polymer physics simulations (Di Pierro et al. 2016). We therefore combined PaleoHi-C data with the OpenMiChroM modeling package (Oliveira Junior, Contessoto, et al. 2021) to construct an ensemble of 3D conformations for woolly mammoth chromosome 8 (Figure S14A). The ensemble exhibits globular structures typically seen for mammalian chromosomes, and yields a simulated contact map (Figure S14B) and contact probability power law (slope = −1.03) consistent with our observations in the woolly mammoth Hi-C data. The structural ensemble can be explored using the Spacewalk genome browser at https://tiny.3dg.io/PaleoHi-C-structures.

To interrogate compartment assignments at individual genes, we generated eigenvector-based A/B annotations in both woolly mammoth and Asian elephant at 500kb resolution. We then looked for bins where the compartment score was strongly discordant in woolly mammoth skin *vs.* Asian elephant skin (Supplementary Online Information). This analysis identified 49 Mammoth Altered Regulation Sequences, dubbed MARS. In aggregate, these MARS span 146 genes (Table S15), including, for example (Figure 3D), *Barx2*, whose deletion in mice leads to short hair (Olson et al. 2005) and *Dusp10*, a regulator of brown adipocyte differentiation (Choi et al. 2013).

### Point-to-point chromatin loops can survive 52,000 years in permafrost

Zooming in further, we explored whether the PaleoHi-C data reflects the presence of point-to-point chromatin loops in the woolly mammoth. Chromatin loops bring pairs of genomic sites that lie far apart along the linear genome into close physical proximity, and are thought to facilitate regulation of genes by distal enhancer elements (Rao et al. 2014). Such loops are thought to form via loop extrusion, a process by which large chromatin loops are generated by first creating a small loop between two adjacent DNA sites, and then sliding these tether points in opposite directions along the 1D genome and thereby increasing the size of the loop (Sanborn et al. 2015; Nichols and Corces 2015; Fudenberg et al. 2016; Darrow et al. 2016). Loop extrusion is known to be mediated by SMC complexes such as cohesin and condensin, and can be arrested by CTCF, leading to the formation of persistent chromatin loops. However, the mammoth contact map is too sparse to observe individual loops, which is typically done using contact maps containing billions of reads.

We therefore looked for evidence of chromatin loops in aggregate. To do so, we generated point-to-point chromatin loop calls using published data from an African elephant fibroblast cell line (Álvarez-González et al. 2022). We used HiCCUPS (Rao et al. 2014) to identify 3578 loops across the African elephant genome (Figure 4, Figure S15). We used these loop calls to perform Aggregate Peak Analysis (APA) (Rao et al. 2014), comparing the aggregate enrichment of the signal from the corresponding positions in woolly mammoth to the enrichment when these positions are translated in any direction (Figure 4). We observed a pronounced enrichment in the center of the APA plot (2.26-fold enrichment relative to the lower-left corner). This enrichment was very similar to the enrichment seen for the same set of loops using in situ Hi-C data from Asian elephant skin (2.18).

**Figure 4.**
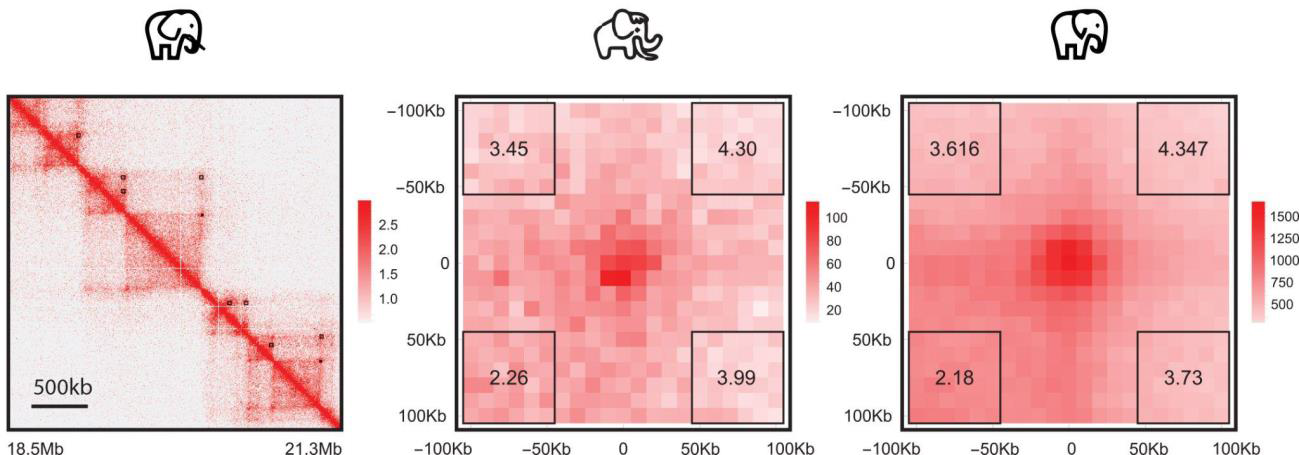
Chromatin loops can survive after 52,000 years in permafrost. **Left**: We called loops in a published Hi-C dataset derived from primary fibroblasts drawn from an African elephant (Álvarez-González et al. 2022). Here we show representative calls from chromosome 28, 18.5Mb - 25.5Mb. Loop calls are indicated using black boxes, above the diagonal; the normalized signal is shown below the diagonal. An interactive version of this panel is available at https://tinyurl.com/2fp789hy. **Center**: Aggregate Peak Analysis of the mammoth data using the African elephant loop list, at 10kb resolution. The central pixel corresponds to summing the 10kb x 10kb region on the Hi-C map surrounding the position of each loop in the African elephant. The intensity of this pixel indicates the number of contacts observed in the woolly mammoth contact map. Other pixels correspond to translating this pixel set a certain number of 10kb intervals in each direction, indicated using the values on the x- and y- axis. Translating this centX and Y axes indicate the number of 10kb bins relative to the position of the loop. The APA score is the ratio of the number of contacts in the central bin to the mean number of contacts in the lower left corner (outlined by the black box in the lower left). Relative enrichments to the other corners are also included. **Right**: A similar APA analysis performed on in situ Hi-C data from modern Asian elephant skin, using the same loop set. The relative signal from loops as compared to the local neighborhood is as strong in the woolly mammoth PaleoHi-C as it is in the modern Asian elephant map. This indicates that chromatin loops can survive for many millennia in permafrost.

These values indicate that many of the loops annotated in the elephant were present at corresponding positions in the woolly mammoth when it was alive, and that, after accounting for the overall reduced frequency of contacts in our ancient sample, loops are as visible in ancient samples as in modern samples. If so, the chromatin proteins underlying loops, such as CTCF and cohesin, may remain relatively intact in the woolly mammoth sample. Furthermore, given enough PaleoHi-C data, it ought to be possible to directly map chromatin loops in woolly mammoth samples without making use of contact mapping data from elephants.

### The inactive X chromosome in woolly mammoth and other elephantids exhibits a tetradic structure

In female eutherian mammals, one of the two X chromosomes is inactivated via a pathway mediated by the noncoding RNA Xist, and exhibits unusual 3D conformations (Brown et al. 1992; Brockdorff et al. 1992). One feature of the inactivated X chromosome (designated Xi) is that it tends to be physically compacted and segregated from the other chromosomes in 3D, manifesting under the microscope as a structure known as the Barr body. This structure is associated with reduced contact frequency between X and other chromosomes that is readily visible in Hi-C contact maps (Rao et al. 2014). Notably, this reduced contact frequency is also apparent in mammoth PaleoHi-C data (Figure 2C), consistent with the preservation of this compacted, segregated Xi conformation for 52,000 years. This observation implies that our sample is female, consistent with the findings of an earlier study based on analysis of aDNA-seq coverage of this sample (Díez-del-Molino et al. 2023).

In humans, the Xi also exhibits two large superdomains, thought to form via extrusion, which partition the chromosome at the CTCF-binding repeat element DXZ4 (Rao et al. 2014; Sanborn et al. 2015; Nichols and Corces 2015; Fudenberg et al. 2016; Darrow et al. 2016). Surprisingly, in the woolly mammoth, we observe a tetradic Xi structure, with four superdomains and three superdomain boundary elements (Figure 5). This tetradic structure is seen in in situ Hi-C maps for female African elephants and female Asian elephants but is absent in their male counterparts (Figure S16). This implies that the tetradic architecture of the inactive X chromosome is conserved in elephantids.

**Figure 5.**
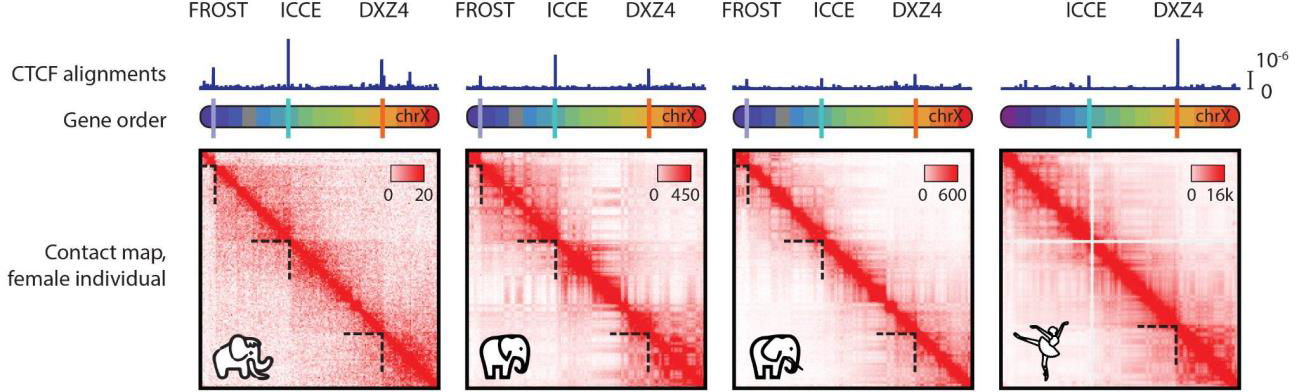
The inactive X chromosome in woolly mammoths and other elephantids exhibits a tetradic structure, with 4 superdomains, distinct from the bipartite structure seen in humans. Contact map for the female mammoth, Asian elephant blood, African elephant primary fibroblasts (Álvarez-González et al. 2022) and human GM12878 B-lymphoblastoid cells (Rao et al. 2014). The chromosomes have been oriented to match the conventional orientation in human. The chromograms above the contact maps indicate the ordering of the loci. The color scheme is based on the human chrX, purple (p-terminus) to red (q-terminus). Corresponding loci across species are shown using the same color. Purple, cyan, and orange ticks on the chromograms indicate the positions of the FROST, ICCE and DXZ4 repeat elements, respectively. The tracks at the top show the number of reads in each map such that at least one of the alignments matches the DXZ4/ICCE-associated CTCF binding motif (Westervelt and Chadwick 2018). Peaks indicate the position of CTCF-binding tandem repeats. Superdomain boundaries are denoted with dashed lines. All tracks are scaled proportionally to the total contact count in the corresponding dataset. The matrices are shown at 1Mb resolution. The Hi-C maps for this figure can be browsed interactively at https://tinyurl.com/2plkschn.

One of the three mammoth Xi boundary elements is DXZ4 (located approximately 200kb proximally from *Pls3* gene in MamPri_Loxafr3.0_assisted_HiC), which is orthologous to the human superdomain boundary.

Another superdomain boundary element is a CTCF-binding tandem repeat called the inactive-X CTCF-binding contact element, ICCE, located ∼35kb downstream of *Nbdy* in MamPri_Loxafr3.0_assisted_HiC (Darrow et al. 2016; Westervelt and Chadwick 2018). Notably, ICCE is present in humans, but does not form a superdomain boundary on human Xi. When we estimated the number of CTCF binding sites associated with the ICCE tandem repeat, we found that, unlike in humans, DXZ4 is not the predominant CTCF-binding locus on the elephantid X chromosome (Figure 5). We hypothesize that when ICCE is longer, it is bound by a larger number of CTCF proteins, making it more effective at arresting extruding SMC-complexes, and giving rise to a superdomain boundary that remains visible after 52,000 years in permafrost.

The third superdomain boundary, which we dubbed FROST, or “Funky Repeat Outlining Superdomain Topology,” lies ∼45kb downstream of *Shroom2* in MamPri_Loxafr3.0_assisted_HiC. Notably, FROST lies at the boundary of the pseudoautosomal region (Figure S16), which is much larger in mammoth and other elephantids as compared to its counterpart in humans (Raudsepp and Chowdhary 2015). To the best of our knowledge, FROST is not orthologous to a tandem repeat that has been annotated in humans.

As expected for structures associated with Xi, the superdomain boundaries at both DXZ4 and ICCE are effaced in samples of male African and Asian elephants (Figure S16). The behavior of the superdomain boundary at FROST cannot yet be resolved in these datasets.

Taken together, our data is consistent with a model whereby long CTCF-binding repeats, which can arrest the extrusion of SMC-family complexes, emerge and disappear on the X chromosome during eutherian evolution, thereby partitioning the Xi (Darrow et al. 2016; Froberg et al. 2018; Westervelt and Chadwick 2018). Of course, we cannot rule out the possibility that other proteins may be responsible, in whole or in part, for the formation of one or more of these superdomain boundaries. Studies examining the effects of modulating ICCE and FROST on Xi architecture and transcription will likely clarify the functional role of these elements.

## Discussion

Over long stretches of time, the topology of DNA (its lengthy chain of intramolecular bonds) is compromised, as intact chromosomes degrade into short fragments typical of ancient samples (Lindahl 1993). Here, we describe a 52,000-year-old sample of woolly mammoth skin that maintains to a high degree, the three-dimensional geometric conformation of the DNA that was present in the cells at time of their death. Thus, our study suggests that the geometry of DNA can survive for far longer than its topology. This may be because a chromosome’s overall shape is reinforced by its protein scaffolding, which remains at least partially preserved in ancient samples. It may also be facilitated by the formation of DNA-DNA crosslinks.

The relatively high degree of conformational fidelity in some ancient samples allowed us to develop PaleoHi-C, which maps chromatin contacts in ancient samples. When used together with a novel algorithm for 3D reference-guided genome assembly, PaleoHi-C data allowed us to generate a reference genome for the woolly mammoth with chromosome-length scaffolds.

Combined with this reference, PaleoHi-C data enabled us to perform detailed epigenetic and nucleomic studies of a 52,000-year-old woolly mammoth skin sample. We find that numerous architectural features seen in modern chromatin samples are preserved after 52,000 years in permafrost. These include qualitative features, such as chromosome territories, the cell-type specific compartmentalization of active (A) and inactive (B) chromatin, domains, loops, and the unique architecture of the inactive X chromosome. They also include quantitative features, such as the power law scalings relating chromatin contact frequency with 1D distance, as well as the relative strength of chromatin loops to the surrounding neighborhood.

Our study yields findings relevant specifically to research on the woolly mammoth, such as revealing detailed gene activation patterns in mammoth as compared to its closest living relative, the Asian elephant. The study also illuminates features of mammalian evolution more broadly, such as the association between dynamic partitioning of the inactive X chromosome and the expansion and contraction of long tandem CTCF-binding repeats.

Overall, our results indicate that for some specimens that are exceptionally well preserved, it may be possible to undertake much more detailed analyses of chromatin architecture, transcriptional activity, structural variation, and even to generate de novo reference genomes. Ultimately extensive testing of the protocol on a wide range of biological materials representative of historic and ancient DNA is needed to fully characterize the preservation conditions in which PaleoHi-C may be applicable.

However, given that (i) like many mammoth skin samples, the skin appeared desiccated at the time of sampling, and (ii) the relatively low rate of cytosine deamination observed, we hypothesize that a critical factor may be rapid desiccation of the sample post mortem. Sample desiccation could occur, for example, through freeze-drying in natural conditions during the extremely cold and dry winters of North-Eastern Siberia, as we suspect applies to this mammoth. (It might also be facilitated for other samples, such as those in natural history or other biological collections, via immediate immersion in ethanol during the sampling process itself.) Given this, we speculate that similarly permafrost preserved samples which are even older than the one described here may be amenable to Hi-C analysis, enabling the study of the evolution of assembled genomes across time, and in multiple Arctic lineages.

Finally, while attempts at species “de-extinction” face formidable hurdles (cf. Lin et al. 2022; Richmond, Sinding, and Gilbert 2016), the potential to generate true de novo reference genomes, to examine the three-dimensional structure of ancient genomes from samples preserved in permafrost, and to explore epigenetic patterns genome-wide may - when combined with developments in the study of ancient RNA (e.g. Fromm et al. 2020; Smith et al. 2019; Fordyce et al. 2013) and proteins (e.g. Cappellini, Collins, and Gilbert 2014; Cappellini et al. 2012) - considerably improve the prospects for success.

## Acknowledgements

We thank Beth Shapiro and Richard E. Green for their advice and guidance during the PaleoHi-C protocol development. We thank Maksim Plikus, Emil Karpinski, George Church, Saul Godinez, Shaiza Pasha, Zane Colaric, Sergei Kliver and Dimoklis Gkountaroulis for insightful suggestions as well as Ishawnia Christopher and SciStories for help with the figures. We thank the animal care and veterinary team at the Houston Zoo for their help with this project. We also thank Dan Fisher for assistance during sampling and J. Camilo Chacón-Duque for his help with the molecular dating analysis.

E.L.A. was supported by the Welch Foundation (Q-1866), a McNair Medical Institute Scholar Award, an NIH Encyclopedia of DNA Elements Mapping Center Award (UM1HG009375), a US-Israel Binational Science Foundation Award (2019276), the Behavioral Plasticity Research Institute (NSF DBI-2021795), NSF Physics Frontiers Center Award (NSF PHY-2019745). E.L.A. and M.A.M.-R. acknowledge support by the National Human Genome Research Institute of the National Institutes of Health under Award Number RM1HG011016-01A1. M.A.M.-R. acknowledges support by the Spanish Ministerio de Ciencia e Innovación (PID2020-115696RB-I00) and the European Research Council under the 7th Framework Program FP7/2007-2013 (ERC grant agreement no. 609989). M.T.P.G., M.S.-V., J.A.R. and C.F. acknowledge the following awards: European Research Council 681396 Extinction Genomics, DNRF143 Center for Evolutionary Hologenomics, and NovoNordisk Foundation NNF21OC0070726 for funding. M.J.R. was supported by the National Institutes of Health grant R35-GM147467. L.D. acknowledges support from the Swedish Research Council (2017-04647 and 2021-00625) and the European Union (ERC, PrimiGenomes, 101054984). J.N.O. is a Cancer Prevention and Research Institute of Texas (CPRIT) Scholar in Cancer Research. J.N.O. also acknowledges NSF (Grant PHY-2210291) and Welch Foundation (Grant C-1792). A.B.O.J. acknowledges the Robert A. Welch Postdoctoral Fellows Program. A.R.-H. acknowledges the Spanish Ministerio de Ciencia e Innovación (PID2020-112557GB-I00) and the Ministry of Economy, Industry and Competitiveness (CGL2017-83802-P). A.L.R. acknowledges: USFWS African Elephant Conservation Fund Grant AFE2129-F22AP01215; UIUC College of ACES Office of International Programs Seed Grant; and Fulbright Denmark Scholar Grant. Genome assembly was performed in association with the DNA Zoo Consortium (www.dnazoo.org). DNA Zoo acknowledges support from Illumina, IBM, and the Pawsey Supercomputing Center. We also thank AMD for the donation of critical hardware and support resources from its HPC Fund.

## Declaration of interests

E.L.A., M.T.P.G. and L.D. are on the scientific advisory board of Colossal Biosciences and hold stock options. M.A.M.-R. receives consulting honoraria from Acuity Spatial Genomics, Inc. E.L.A. and O.D. are inventors on US provisional patent application 16/308,386, filed 7 December 2018, by the Baylor College of Medicine and the Broad Institute, relating to the assembly methods in this manuscript. E.L.A. and O.D. are inventors on US provisional patent application 62/617,289, filed 14 January 2018, and US provisional patent application 16/247,502, filed 14 January 2019, by the Baylor College of Medicine and the Broad Institute, relating to the assembly methods in this manuscript. E.L.A. is an inventor on US provisional patent application PCT/US2020/064704 filed 11 December 2020, by the Baylor College of Medicine and the Broad Institute, relating to the assembly methods in this manuscript.

## Data availability

Raw FASTQ data for the woolly mammoth can be downloaded from https://tiny.3dg.io/Globe-KU-PaleoHi-C. The modern Asian elephant and domestic donkey Hi-C data are available via Sequencing Read Archive (SRR11097134, SRR11097135, SRR11097167, SRR11097169, SRR11097170, SRR25023864-SRR25023867 and SRX5415918, SRX5415921, respectively). Fastas and annotations are available at https://tiny.3dg.io/PaleoHi-C-fastas.

## Author Contributions

M.T.P.G. and E.L.A. jointly conceived of this project. M.A.M.-R., M.T.P.G. and E.L.A. oversaw data analysis. V.P. and L.D. provided woolly mammoth tissue samples. I.G.-T., R.C., J.P.F. and K.P. provided elephant samples. M.S.-V. and C.P.E. developed ancient Hi-C protocols, and performed Hi-C experiments on ancient samples with technical assistance from S.S.T.M. O.D., R.K. and A.D.O. performed Hi-C experiments on modern samples. O.D. led the development of genome assembly algorithms and performed genome assembly, with assistance from R.K., D.W., S.S.B., M.S.S., J.A.R., M.A.M.-R., N.C.D., and A.R.-H. O.D., J.A.R., M.D., and C.F. performed bioinformatics analyses, with input from A.K. and M.J.R. (CRUSH). V.G.C. and A.B.O.J. performed polymer physics simulations under the supervision of J.N.O. M.S.-V., O.D., C.P.E., S.S.T.M., B.O., A.G., E.S.L., M.T.P.G., and E.L.A. contributed to PaleoHi-C protocol development. O.D., J.A.R., M.S.-V., A.L.R., M.A.M.-R., M.T.P.G. and E.L.A. wrote the manuscript with input from all authors.

