## Supplementary Online Information for "Three-dimensional genome architecture persists in a 52,000-year-old woolly mammoth skin sample"

**Sample collection and handling details.**

The specimen (ID IN18-032) was collected in September 2018 near Belaya Gora, Sakha Republic (N68.57887, E147.16055) during an international expedition. It comprised a fragment of a skin from a woolly mammoth head (see Figure S1). Upon visual inspection, the skin appeared to be very well preserved, with an intact ear and no signs of putrefaction.

Two fragments from the rim of the skin sample from behind the ear were collected for research purposes several hours after the sample was extracted from permafrost. One was kept in 95% ethanol and used for PaleoHi-C experiments as well as for generating the shotgun Illumina data (aDNA-Seq) published in (Díez-del-Molino et al. 2023). A second larger fragment was not placed in ethanol, and was transported 'as is' until it could be stored frozen for longer term storage. This sample was intended for radiocarbon dating, and the leftovers have been used for supplementary PaleoHi-C experiments. A closeup photo (Figure S1A) shows the second (no-ethanol) fragment. The ethanol sample was collected in the immediate vicinity of the one in the closeup as seen in Figure S1D.

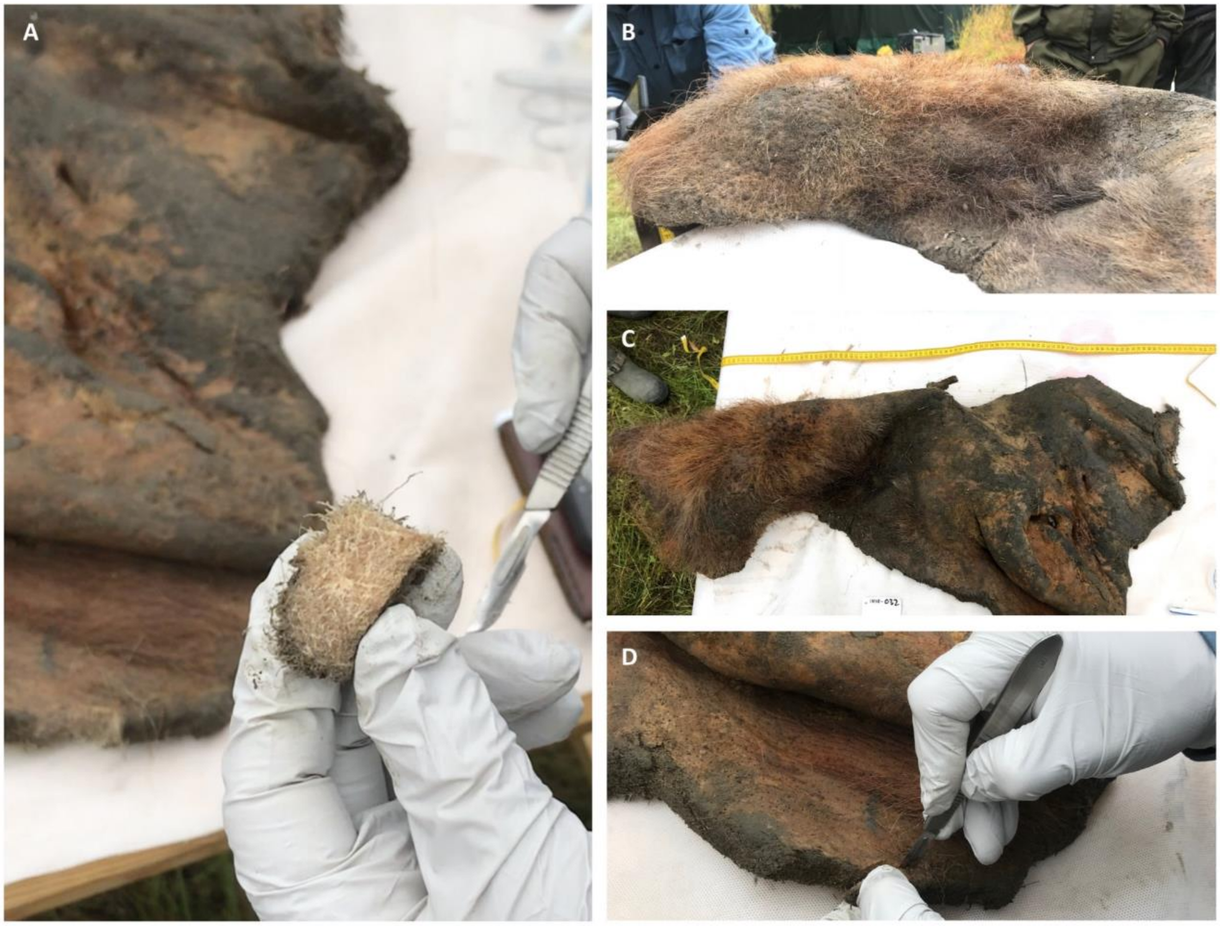

*Figure S1. (A-D): Photographs of the mammoth sample used in this study (specimen ID IN18-032) showing good preservation and no signs of putrefaction. A close-up in (A) shows one of the two samples that have been collected for research purposes. A rim of the skin from behind the ear from where the two samples were cut, side by side, is shown in (D).*

**Radiocarbon dating.**

The sample was sent for radiocarbon dating to the Oxford Radiocarbon Accelerator Unit (ORAU), which yielded an infinite age estimate (>44,900, OxA-38763) (Díez-del-Molino et al. 2023).

**Phylogenetic dating.**

To obtain a more precise estimate of the sample's age, we used a Bayesian molecular dating method based on the mitochondrial genome following the approach implemented in (van der Valk et al. 2021). First, using the published aDNA-Seq data from (Díez-del-Molino et al. 2023) the mitogenome was assembled for IN18-032 from merged and trimmed FASTQ files containing DNA fragments over 35 base pairs using mapping-iterative-assembler (MIA) (Green et al. 2010) with the Asian elephant mitogenome (NC_005129, (Rogaev et al. 2006)) as a reference. Given the vast amount of whole-genome sequencing data, only one library (accession: ERR10173648) was used for this analysis. This resulted in a mitogenome coverage of 800x. Positions with less than 3x coverage or with a sequence agreement of less than 67% were assigned as missing data.

Next, the assembled mitogenome was aligned to previously published elephantid mitogenomes compiled by van der Valk *et al*. 2021 with Muscle v3.8.31 (Edgar 2004). The obtained multiple sequence alignment (containing a total 172 mammoths, nine African elephants, two Asian elephants and four mastodons) was visually inspected using SeaView v5.0.5 (Gouy et al. 2021). Considering the assembly and alignment limitations on the hypervariable control region, the VNTR region as identified in the woolly mammoth reference mitogenome (NC_007596 (Krause et al. 2006), positions 16157 to 16476) was removed from the alignment. Additionally, sequences supported by only one sample in the control region (and assigned as gaps in the remaining samples) were also removed.

Next, we performed a Bayesian phylogenetic analysis using BEAST v1.10.4 (Suchard et al. 2018). In order to avoid over-parameterising the model for tip dating and to reduce the uncertainty of the age estimate (Chacón-Duque *et al*., in prep), we removed all samples without finite calibrated radiocarbon ages from the alignment, in order to obtain a tip date estimate only for the sample of interest. This filtering removed all mastodons and left a total of 121 radiocarbon-dated mammoths. Following van der Valk *et al*. 2021, the alignment was split into six partitions: tRNA, rRNA, first, second, and third codon positions, and the control region and all were assigned the HKY + Gamma + Invariant substitution model, except for the tRNA partition, for which the HKY + Invariant model was used instead (van der Valk et al. 2021). Also, the same log-normal prior for the divergence between *Loxodonta* and *Elephas/Mammuthus* of 5.3 Ma was used with a strict molecular clock and a flexible skygrid coalescent model (Gill et al. 2013). Two Markov Chain Monte Carlo (MCMC) chains using a uniform tip prior (date: 75 ka, range: 10 ka - 2 Ma) were run for 100,000,000 iterations, sampling each 10,000 and discarding the first 10% as burn-in. We checked convergence with Tracer v1.7.2 and combined the chains with LogCombiner v1.10.4 (Suchard et al. 2018). This resulted in a final age estimate of 52.3 ka (95% HPD: 14.4-86.9 ka).

**PaleoHi-C protocol.**

*Sample preparation, crosslinking and aDNA-chromatin extraction*. The sample was divided into 80 mg subsamples, which were first coarsely crushed using a scalpel. 500µl of 1% formaldehyde was added as crosslinking solution, and the sample was left to incubate at room temperature (RT) for 10 minutes in slow rotation. 40µl of 2.5M glycine solution was added to the reaction, vortexed and incubated for 5 min at RT. Sample was centrifuged for 2min at 6000 x g and the supernatant was discarded into appropriate chemical waste. 500µl of Wash Buffer1 (10mM Tris-HCl pH 8, 50mM NaCl and 0.01% Tween) was added and mixed to wash. Sample was centrifuged for 2 min at 6000 x g and the supernatant was discarded. The wash was repeated for a total of two washes.

After washing the crosslinked tissue, the sample was grinded using a plastic pestle. While keeping the sample the sample in a cold rack, 550µl of freshly prepared Lysis buffer (10mM HEPES pH 8, 10mM NaCl, 0.2% IGEPAL 630, 1X Protease inhibitors) were added and the reaction was vortexed thoroughly. The reaction was left to incubate in a cold rack inside the fridge for 30 min. The reaction was vortexed again and homogenized with a plastic pestle with 30 up-and-down strokes inside the tube. Samples were spun at 2500 x g for 5 min at RT. (The supernatant from this step was collected to prepare supplementary datasets, see *Detailed description of generated datasets*.) 500µL of Wash Buffer2 (50mM Tris-HCl pH 8, 50mM NaCl, 1mM EDTA) were added without disturbing the tissue pellet. Sample was spun at 2500 x g for 5 min at RT to re-collapse the tissue pellet between washes and supernatant was discarded. The sample was washed a second time with 500µL of Wash Buffer2, spun at 2500 x g for 5 min at RT and supernatant was discarded. Sample was resuspended in 250µL of Wash Buffer2 and 100µL of 2% SDS were added. The reaction was vortexed to mix everything thoroughly and left to incubate with shaking for 10 min at 50°C. After incubation, the supernatant containing crosslinked aDNA-chromatin was collected and an estimate of DNA recovered made using a Qubit instrument with dsDNA HS assay.

*Bead binding, chromatin digestion, end repair and biotin incorporation, end ligation*. Carboxylated beads (2X) were added to the supernatant collected previously. The reaction was mixed by pipetting 10 times up and down. (Different beads such as SPRI, AmpureXP and Magbio beads were used to bind aDNA-chromatin in solution from different tissue replicates and no significant correlation between library quality and bead type was found, see Table S1.)

The reaction was incubated for 5 min at RT. The mix was placed on a magnet for 5 min, and the supernatant was discarded. The bead-bound samples were then washed with 500µL of Wash Buffer1. Again, the supernatant was discarded and the beads were washed a second time with 250µL of Wash Buffer1.

After removing the supernatant from the last wash, 70µL of restriction enzyme digestion mix containing 5U of DpnII was added to the beads and the reaction was left incubating at 1000 rpm for 1 hr at 37°C. Sample was then placed on a magnet for 5 min, the supernatant was discarded and the beads were washed twice with 500µL of Wash Buffer1.

Next, 50µL of a reaction mix containing 10X NEBuffer 2, 10mM of each dATP, dTTP and dGTP, 0.4mM Biotin-14-dCTP and 5U/µL Klenow Polymerase Large Fragment was added to each sample and the reaction was left to incubate at 1000 rpm for 30 min at 25°C. Afterwards the reaction was placed on a magnet, supernatant was discarded and the bead-bound sample was washed twice with Wash Buffer1. Finally, 500µL of ligation reaction mix containing 10X T4 DNA ligase buffer, 20 mg/mL BSA, 10% Triton X-100 and T4 DNA Ligase 400 U/µL was added to each sample, and reactions were left to incubate at 500 rpm for 2 h at RT.

*Nucleotide exchange, crosslink reversal and DNA purification.* While keeping the tubes on a thermoblock, 2.5µL of 10mM of dNTP mix and 2.5µL of T4 DNA polymerase were added to each sample, and the reactions were left incubating further at 1000 rpm for 15 min at 16°C. Samples were then placed on a magnet, supernatant was discarded, and samples were washed once with 250µL Wash Buffer1. Next, a crosslink reversal reaction to separate DNA from proteins and degrade the leftover proteins was carried out by adding 100µL of CRMix containing 50mM Tris-HCl pH 8, 1% SDS, 0.25mM $CaCl_2$ and 1mg/ml Proteinase K. The reactions were left to incubate at 1000 rpm for 15 min at 55°C followed by 45 min at 68°C. After incubation, samples were placed on a magnet, and the supernatant was transferred to new 1.5mL Eppendorf tubes. 200µL of carboxylated beads (SPRI, AmpureXP or Magbio) were added, mixed by pipetting. After incubating for 5 min at RT samples were placed on a magnet, the supernatant was discarded, and the bead-bound samples were washed with 250µL of freshly prepared 80% EtOH while still on a magnet. EtOH was removed and samples were washed again for a total of 2 washes. After the second wash, samples were left to air-dry for 5 min. 58µL of EB Buffer were added, and samples were incubated for 5 min at 37°C.

*Library preparation, cleanup and amplification.* Purified DNA was prepared for sequencing on two platforms, Illumina and BGISEQ. 27µL for each library were aliquoted and libraries were prepared following the BEST protocol (Carøe et al. 2018). After the fill-in step, samples were cleaned using C1 Streptavidin beads. 60µL of C1 beads per sample were washed and resuspended in 130µL of 2X NTB Buffer (10mM Tris-HCl, 2M NaCl, 1mM EDTA). The mix was then added to each sample and the reaction was left to incubate at RT for 30 min with slow movement (300 rpm). After incubation, samples were placed on a magnet, supernatant was discarded and the beads were washed twice with 200µL of Buffer NWB (10 mM Tris-HCl, 1M NaCl, 1mM EDTA, 0.05% Tween 20) followed by two washes with 200µL of TWB buffer (10mM Tris-HCl, 0.5mM EDTA, 0.05% Tween 20). While on a magnet, supernatant was removed, and beads were resuspended in 94µL of Amplification Master Mix (1X Kapa U+ HotStart ReadyMix, 10 mg/mL BSA). The mix was split into two 47µL reactions that were each supplemented with 0.3uM of each Fwd. and Rev. indexes to a final volume of 50µL per reaction. The replicates were PCR-amplified for 16-20 cycles. The reactions were then cleaned using 1.2X SPRI beads and eluted into a final volume of 32µL with EB Buffer. Following quantification and visualization of amplified PaleoHi-C libraries on an Agilent TapeStation instrument, samples were submitted to BGI Europe for either BGISEQ or Illumina sequencing.

**Detailed description of generated datasets.**

Overall, 26 mammoth Hi-C datasets were generated: six "canonical" PaleoHi-C libraries from the ethanol-preserved sample and 20 supplementary datasets.

The "canonical" datasets were generated as follows: the ethanol-preserved tissue was split into two biological replicates (~80mg each). The PaleoHi-C protocol was carried out on these as described in the PaleoHi-C protocol section until the step when crosslinks were reversed and the DNA purified. The resulting material has been split in two, with one half prepared for downstream sequencing on the Illumina platform, resulting in two libraries (one library from each of the tissue replicates), and the other half for the BGI sequencing platform. The material being prepared for BGI sequencing has been further split before the amplification step and processed as two separate PCR reactions, resulting in four libraries, two for each biological tissue replicate. The resulting datasets are designated as "EtOH precipitate" in Figure S2 and Table S1.

Supernatant material from the tissue lysis step (see *PaleoHi-C protocol* section) was collected and taken though the steps downstream of lysis, as described in the protocol section, to convert into a set of supplementary libraries. Similar to the "canonical" set, the material was split to be prepared for the Illumina and BGI sequencing platforms after DNA purification. DNA was split in two for running PCR amplification when preparing the BGI libraries. This resulted in six "EtOH supernatant" supplementary datasets.

We additionally explored the viability of the PaleoHi-C method on leftovers of the second skin sample that had been collected for 14C dating in the field, thus not preserved in ethanol. Four replicate subsamples were taken from this sample, and the PaleoHi-C protocol was carried out as described above, starting with the crosslinking step. Purified DNA was prepared for sequencing on the BGI sequencing platform. All reactions but one were split before the amplification step and processed as two separate libraries. The procedure resulted in seven supplementary datasets (designated "No EtOH precipitate").

Finally, post tissue-lysis supernatant material from the no-ethanol sample experiments was converted into “No EtOH supernatant” supplementary libraries. Just as with the corresponding precipitate datasets, purified DNA was prepared for sequencing on the BGI sequencing platform. All reactions but one were split before the amplification step and processed as two separate libraries.

Detailed statistics associated with all 26 datasets, as generated by the Juicer Hi-C processing pipeline (Durand et al. 2016) can be found in Table S1 (Juicer version 1.6; BWA 0.7.17-r1188; 8 threads; openjdk version "1.8.0_222-ea"; Juicer Tools Version 1.9.9). Data was aligned to the African elephant genome reference Loxafr3.0 (GCA_000001905.1, (Palkopoulou et al. 2018)), and the pipeline was run with “-s none” flag to account for non-enzymatic fragmentation of the sample. The distribution of the percent of total alignable read pairs in the canonical and supplementary datasets is shown in Figure S2.

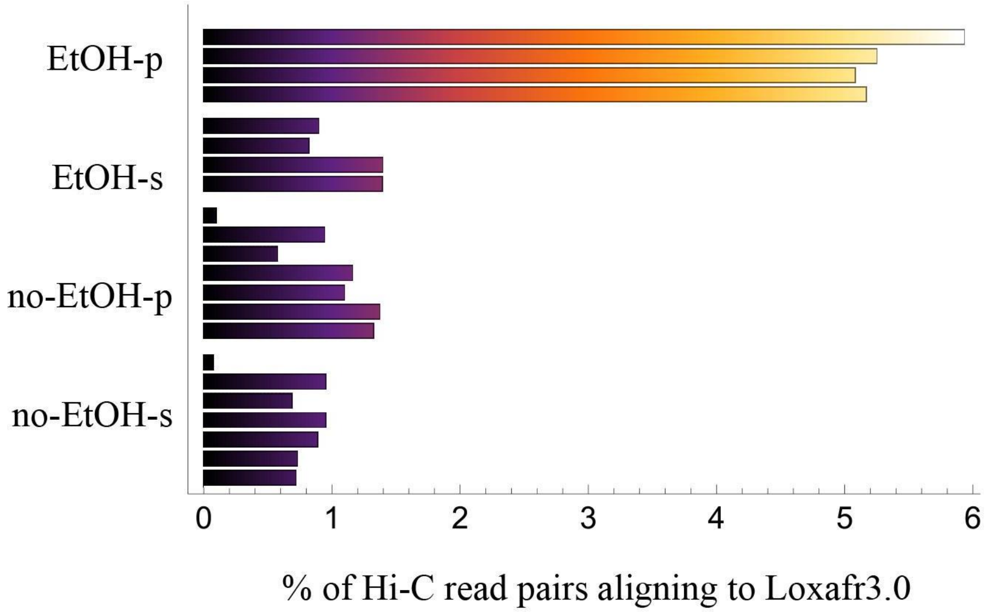


*Figure S2. Immediate on-site preservation of sample in ethanol prevents microbial activity and allows isolation of more endogenous mammoth Hi-C sequences when compared to a sample handled without a preservative. Although with lower efficiency, endogenous contact data can be collected from non-ethanol samples as well. Percent of total alignable read pairs (calculated with respect to the total raw read pair count) are listed as reported by the Juicer pipeline (Rao et al. 2014; Durand et al. 2016). Alignment statistics for 22 BGI libraries are shown. The four Illumina libraries are omitted to avoid collating the systematic bias introduced by the sequencing platform from the signal intrinsic to the datasets. The libraries are bundled into four groups: EtOH-p for “EtOH precipitate”, the main PaleoHi-C dataset; EtOH-s for “EtOH supernatant” supplementary dataset prepared from the tissue lysis supernatant leftovers; no-EtOH-p for the “No EtOH precipitate” supplementary dataset generated from the sample handled without ethanol; and no-EtOH-s for the “No EtOH supernatant” supplementary dataset generated from the supernatant collected during no-ethanol tissue lysis. The datasets are arranged from top to bottom in agreement with the order in which they appear in Table S1, left to right.*

**Modern elephant samples and library preparation.**

We generated in situ Hi-C data for five different Asian elephant tissues: skin, liver, ovary, brain and peripheral blood mononuclear cells (PBMCs). The skin, liver, ovary and brain tissue samples were collected during a necropsy of a single female Asian elephant individual (Elephas maximus indicus) performed in 2004 at the San Antonio Zoo. The in situ Hi-C libraries, one for each tissue, were prepared as described in (Rao et al. 2014), and a combination of Csp6I and MseI restriction enzymes was used for chromatin digestion.

We generated PBMC in situ Hi-C data from two different Asian elephant individuals, one female and one male. Ear vein blood samples for both preparations were opportunistically collected during routine veterinary procedures at the Houston Zoo. Two in situ Hi-C libraries were prepared as described in (Rao et al. 2014), and MboI restriction enzyme was used for chromatin digestion in both cases.

All samples were obtained under Baylor College of Medicine protocol AN-6832. The in situ Hi-C libraries were sequenced on the Illumina sequencing platform.

**Non-USER treated aDNA-Seq.**

An extensive aDNA-Seq dataset for sample IN18-032 was published in (Díez-del-Molino et al. 2023). We supplemented the published sequencing reads with a small additional dataset.

Just as in the case of data published in (Díez-del-Molino et al. 2023), DNA was extracted from a piece of the ethanol preserved tissue sample at the ancient DNA lab at the Swedish Museum of Natural History, Stockholm, Sweden. The tissue was digested overnight with the lysis buffer described in (Sinding et al. 2015). Next, the lysis was concentrated and purified as described in the supplementary protocol of (Dehasque et al. 2022). Double-stranded sequencing library was prepared following the protocol of (Meyer and Kircher 2010) with one exception. Specifically, we omitted the USER treatment during the blunt-end repair step to excise uracil bases from the DNA fragments as described in (Pečnerová et al. 2017). The libraries were sequenced using the Illumina platform and sequencing data was aligned to Loxafr3.0 using Juicer version 1.6/BWA 0.7.17-r1188 (Durand et al. 2016). Alignment statistics can be found in Table S3.

**PaleoHi-C contact probability as a function of genomic separation.**

In order to analyze the decay of read separation probability with genomic distance in aDNA-Seq, PaleoHi-C experiments and in situ Hi-C data generated using a modern Asian elephant skin sample, we aligned all datasets to Loxafr3.0 (GCA_000001905.1, (Palkopoulou et al. 2018)) using Juicer (version "1.6", (Durand et al. 2016)). The pipeline was run with "-s none" flag for all datasets and, in the case of aDNA-Seq data, with an additional “-j” flag (stands for “just exact duplicates excluded at the deduplication step”) supplied for handling very large low complexity datasets.

Only high-quality alignments (mapping quality greater than or equal to 30 for both sequences in the read pair) to the largest scaffold in Loxafr3.0 (GL010027.1) were extracted for the contact probability analysis. Logarithmic binning was used to smooth the curves. Values have been normalized by the total number of contacts within GL010027.1 in each corresponding experiment.

All other things equal it would carry greater significance to compare the PaleoHi-C data (not treated with the USER cocktail) to the non-USER-treated aDNA-Seq library for the same sample since exposure to the USER enzyme cocktail has the capacity to reduce the DNA fragment length. In practice, the final size distribution depends on other experimental factors such as, e.g., DNA quality variation, amplification conditions and cleanup procedure details. In view of this, we examined the size distribution of the treated and untreated aDNA-Seq datasets and chose the USER-treated dataset from (Díez-del-Molino et al. 2023) for the most conservative comparison with PaleoHi-C 1D separation probability distribution (see Figure S3A). Only a subset (ERR10173641) of a full dataset was used for Figure 1D. The full dataset is included in Figure S3, panel A.

The Illumina-sequenced PaleoHi-C dataset has been used when comparing to the Illumina shotgun aDNA-Seq data in Figure 1D to avoid possible bias associated with a particular sequencing platform. Figure S3A includes a version of the same plot with PaleoHi-C data curve generated from data collated from across the BGI and Illumina sequencing platforms.

In order to provide further insights, we include two additional supplementary figures. Figure S3B compares the mammoth skin PaleoHi-C dataset with the modern Asian elephant skin sample across a wider range of distances. Figure S3C presents a breakdown of contact probability curves across the four dataset groups described in *Detailed description of generated datasets* (EtOH precipitate, EtOH supernatant, no-EtOH precipitate and no-EtOH supernatant). All datasets show the characteristic contact probability decline, with approximately the same slope. Note however the “noise” associated with the curves aside from that corresponding to the EtOH-p dataset (the ‘canonical’ PaleoHi-C) on account of relatively little data generated by these protocol variants. The analysis of the variants is done using BGI sequencing data.

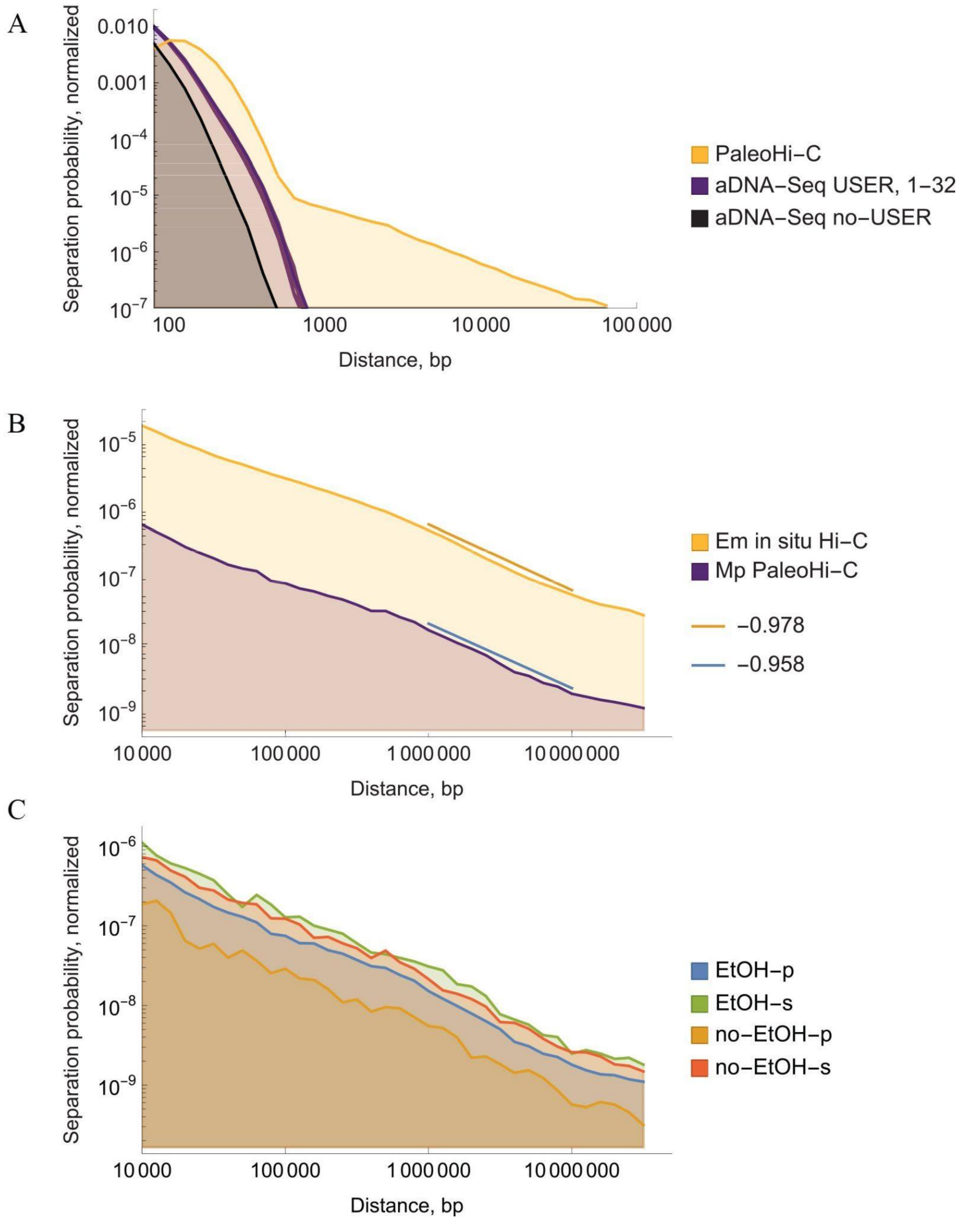


*Figure S3. Supplementary details on the probability of observing a read pair in which the two reads map to positions in the genome separated by a given distance in basepairs. (A) A small fraction of PaleoHi-C data reflects contacts between loci that lie far away in 1D. No such fraction exists in aDNA-Seq datasets. The image is analogous to that*

*shown in Figure 1D, with the following modifications: 1) the PaleoHi-C curve is based on data collated from both the BGI and Illumina sequencing platforms; 2) two aDNA-Seq datasets are included: the USER-treated datasets (from across 32 replicates) from (Díez-del-Molino et al. 2023) and the non-USER treated dataset generated for the same sample as part of this study. The more contiguous dataset from (Díez-del-Molino et al. 2023) was chosen to be plotted alongside PaleoHi-C data in Figure 1D for a more conservative comparison. (B) Contact probability curves for PaleoHi-C of woolly mammoth skin and in situ Hi-C of Asian elephant skin. The slopes between the two datasets are highly consistent across a wide range of distances. (C) Contact probability curves plotted separately for the "main" PaleoHi-C dataset (EtOH-p for EtOH precipitate dataset) and the 3 supplementary datasets: EtOH-S (EtOH supernatant), no-EtOH-p (no-EtOH precipitate) and no-EtOH-s (no-EtOH supernatant).*

#### DNA damage in PaleoHi-C data.

DNA damage is well documented in ancient and degraded DNA samples, and is most frequently evident in DNA sequence data in the form of cytosine (C) deamination (Lindahl 1993; Pääbo 1989; Hansen et al. 2001), which are then converted to uracil (U) and its analogues, thus revealed as cytosine to thymine (T) transitions in resulting sequence data.

To find the proportion of PaleoHi-C reads bearing this damage pattern we used PMDtools (Skoglund et al. 2014), https://github.com/pontussk/PMDtools. The software assigns each read a score (PMDscore) for which positive values indicate post-mortem damage, with the higher scores associated with higher degrees of damage. The code was run with default parameters to compute the percentage of damaged reads in generated datasets (reads with PMDscore>3), using PaleoHi-C data alignments to Loxafr3.0.

In addition to the PaleoHi-C dataset we analyzed one of the in situ Hi-C libraries for the Asian elephant (skin necropsy sample) and the African elephant in situ Hi-C dataset from (Álvarez-González et al. 2022), SRR19650826. We also computed the percentage of reads with damage in a USER-treated shotgun whole genome sequencing (aDNA-Seq) experiment for the same mammoth individual published in (Díez-del-Molino et al. 2023) and the non-USER treated aDNA-Seq dataset.

The mammoth-derived PaleoHi-C reads have an elevated degradation signature, on par with the non-USER treated aDNA-Seq data (Figure S4A and Table S4). The damage is higher than in any of the modern elephant datasets. The result remained true when analysis was run specifically for long-range 3D contacts (i.e. using only reads that represent interactions between loci occurring on the same Loxafr3.0 scaffold and separated by 1D distance greater than 20kb).

#### Authentication of mammoth reads using diagnostic positions.

To ascertain the authenticity of our mammoth sample, we run a diagnostic test using a set of single nucleotide polymorphism (SNPs) to discard the unlikely scenario of contamination from modern closely related *Elephantidae* species in our mammoth PaleoHi-C sample.

For this, we first obtained mammoth diagnostic positions from a SNP panel from a recent study (Díez-del-Molino et al. 2023) of 29 elephants (7 Asian elephants and 21 African elephants) and 23 woolly mammoths using VCFtools (Danecek et al. 2011). We consider a site to be diagnostic when all elephants have the reference allele, and the mammoths always show the alternative allele, and requiring that at least 4 mammoths and 14 elephants have data for the position. In total, we used 1,034,282 diagnostic positions.

We then proceeded to analyze the alignments at the diagnostic positions in the PaleoHi-C dataset. In addition to the PaleoHi-C dataset we analyzed one of the in situ Hi-C libraries generated for the Asian elephant (skin necropsy sample) and the African elephant in situ Hi-C dataset from (Álvarez-González et al. 2022), SRR19650826.

Together with our mammoth Hi-C data, all datasets were mapped to the African elephant genome assembly Loxafr3.0 (Palkopoulou et al. 2018) in a single-end mode to recover as many reads as possible. We applied bwa-0.7.17 mem for mapping the African and Asian elephant Hi-C libraries and bwa aln with parameters "-l 16500 -n 0.01 -o 2" for mapping the mammoth Illumina and BGI PaleoHi-C (Li and Durbin 2010; Li 2013). We removed duplicates for each library with picard v2.15.0 (http://broadinstitute.github.io/picard/) MarkDuplicates. BAM files from the same sample were merged with picard v2.15.0 MergeSamFiles. We filtered for primary alignments using samtools view -F 260 (Danecek et al. 2021).

To account for variable coverage depth between different datasets, we performed haploid calling by randomly sampling one base at each diagnostic position using ANGSD v0.940 (Korneliussen, Albrechtsen, and Nielsen 2014) with the following parameters: "-doHaploCall 1 -doCounts -rf positions_mammoth_fixed_angsd.bed -uniqueOnly 1 -remove_bads 1 -minMapQ 30". We then counted the total covered sites per sample and the positions showing the mammoth or the elephant allele.

We found that reads from mammoth libraries - when they could be assigned - were overwhelmingly assigned to mammoth (99.8% of the time) and almost never assigned to elephant (0.2%), whereas reads from elephant libraries were overwhelmingly assigned to elephant (99.9%) and almost never assigned to mammoth (0.1%). The same result can be obtained using only a subset of reads that represent long-range 3D contacts for the mammoth and modern elephants, see Figure S4B and Table S5. The few cases of discordant assignments could be explained by sequencing errors. We conclude that the signal in PaleoHi-C experiments derives from ancient mammoth DNA, rather than from modern contaminants.

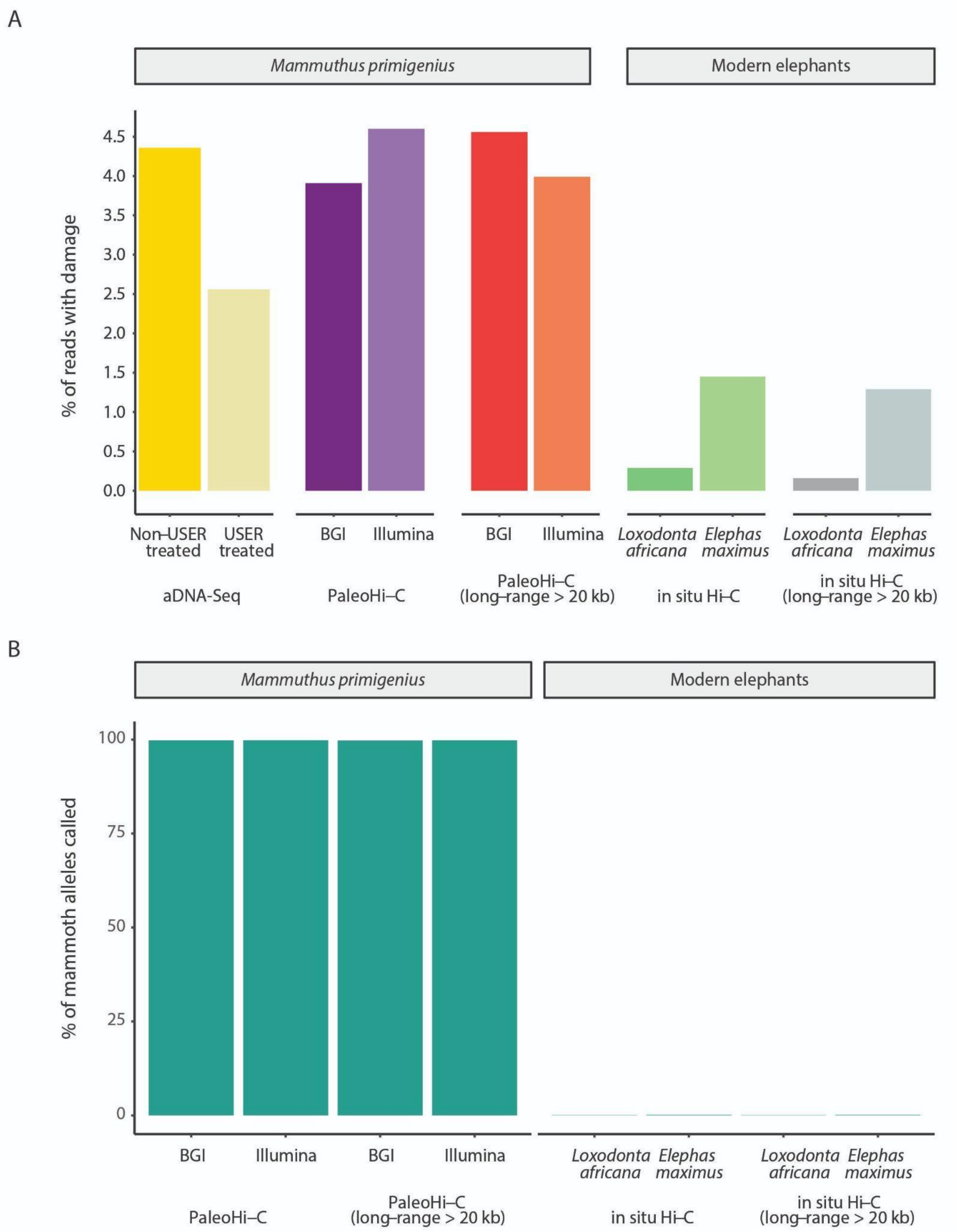


*Figure S4. A. Barplot showing the percentage of reads that display damage related to cytosine deamination in the woolly mammoth sample and the modern elephant samples, as determined by PMDtools (Skoglund et al. 2014). As*

*expected, the mammoth-derived PaleoHi-C reads have an elevated degradation signature, on par with the non-USER treated aDNA-Seq data. The damage is higher than in any of the modern elephant datasets. Note the elevated damage signature in the skin necropsy for the Asian elephant (*Elephas maximus*) as compared to the fibroblast-derived African elephant (*Loxodonta africana*) dataset. The relative increase in percent of damaged reads is consistent with the necropsy sample being stored under suboptimal conditions for ~20 years. B. Barplot showing the percentage of mammoth alleles in reads overlapping mammoth-specific fixed genetic variants from (Díez-del-Molino et al. 2023), in PaleoHi-C data vs. data derived from modern elephants. Whenever such diagnostic reads are examined in the mammoth, they overwhelmingly show the mammoth allele, confirming that PaleoHi-C data derives from ancient mammoth DNA rather than from modern contaminants.*

**Domestic donkey Hi-C data.**

The donkey Hi-C data was generated from peripheral blood mononuclear cells that were extracted from a blood sample bought from Animal Technologies, Inc. (https://www.animaltechnologies.com/). Two replicate libraries were prepared following the in situ Hi-C protocol described in (Rao et al. 2014). MboI restriction enzyme was used for chromatin digestion. The replicate libraries were sequenced on the Illumina HiSeq 2000 instrument, and data uploaded to the Sequencing Read Archive (SRA data accession SRX5415918 and SRX5415921).

**Domestic donkey assisted genome assembly.**

The donkey in situ Hi-C sequencing data was aligned to the EquCab2.0 (GCF_000002305.2, (Wade et al. 2009)) reference and deduplicated using Juicer version 2.0/BWA 0.7.17-r1188 (Durand et al. 2016). The pipeline was run with “-s MboI” flag. The resulting statistics for both libraries as generated by the Juicer pipeline are listed in Table S6.

The resulting alignments were used as input to the 3D-DNA pipeline, v201008 (Dudchenko et al. 2017). The pipeline was run with default parameters to identify chromosomal breaks, fusions and intrachromosomal rearrangements that separate the donkey and the horse and “edit” the horse reference to reflect the said events. The resulting candidate assembly representing a tentative donkey karyotype was examined and polished using Juicebox Assembly Tools (Dudchenko et al. 2018). The Hi-C data aligned to the original EquCab2.0 reference and the “donkified” EquCab2.0 are shown in Figure S5.

The core steps of the 3D-DNA pipeline are: 1) misjoin detection; and 2) anchoring, ordering and orientation (scaffolding) of the resulting reference fragments. When performing an assisted assembly from a high-quality reference such as EquCab2.0 the former step primarily detects evolutionary breaks which, just as “true” misjoins, appear as a disagreement between the Hi-C data and the underlying reference in the form of a near-diagonal depletion of Hi-C proximity signal not typically observed in contact maps of truly contiguous fragments. Similarly, the second step primarily detects evolutionary fusions that appear as increased Hi-C contacts between different reference fragments. Of course, in the case of an imperfect or incomplete reference assembly the same steps would detect misjoins and scaffold draft sequences just as they would as part of a de novo Hi-C-guided genome assembly workflow.

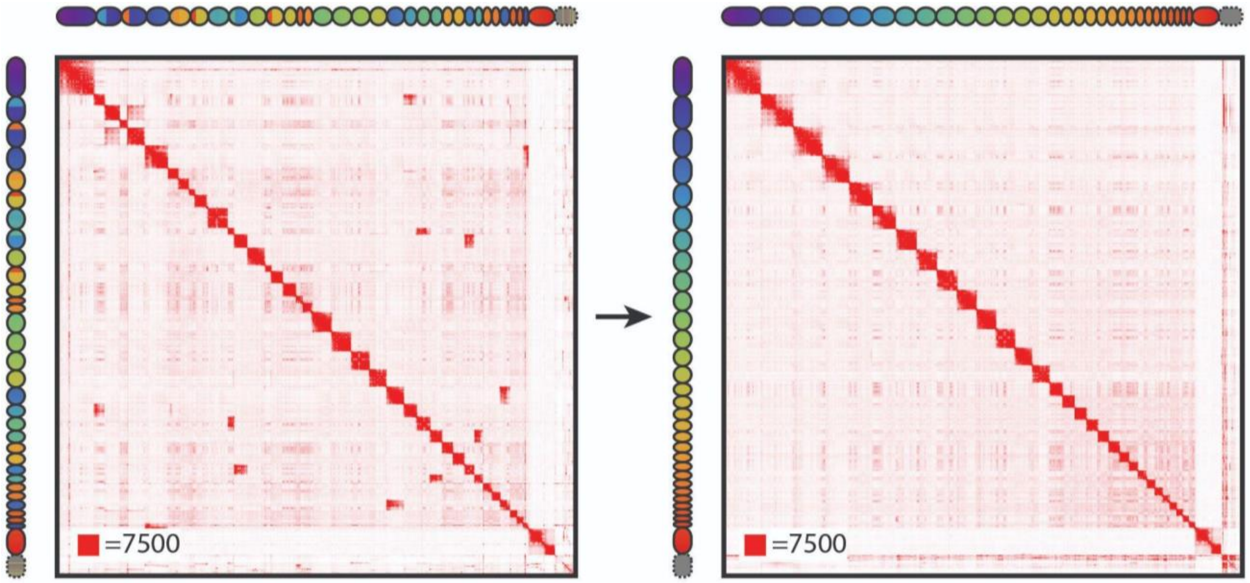


*Figure S5. Starting with the EquCab2.0 domestic horse genome assembly (2n=64, (Wade et al. 2009)), left, we used Hi-C data for the donkey* Equus asinus *to identify the evolutionary events such as chromosome breaks, fusions as well as intrachromosomal rearrangements, that separate the two species. The events manifest as near-diagonal depletions and off-diagonal enrichments that suggest a disagreement between the 1D distance separating a pair of genomic loci as suggested by the Hi-C data and by the underlying reference. (Simplifying somewhat, off-diagonal enrichments suggest that the sequences are further away on the reference than they should be according to Hi-C data, and vice versa, a depletion near diagonal that is not associated with a reference sequence boundary suggests that the sequences flanking the depletion should be moved further away from each other along the 1D reference.) We then proceeded with editing the horse chromosomes, introducing breaks and fusing the horse sequences to reflect the identified evolutionary events, until the Hi-C data and the underlying edited reference are in agreement, with the resulting map showing no off-diagonal enrichments or near-diagonal depletions not associated with chromosome boundaries, on the right. The resulting reference is a close approximation of the donkey genome (2n=62). The contact maps show the same donkey Hi-C data, aligned to the original EquCab2.0 and lifted over to the rearranged reference. Rainbow tracks on top and to the left of the contact maps are used to highlight synthetic sequences between the two references: the same color is used to show corresponding loci. The ovals shaping the rainbow tracks outline the boundaries of the 32 chromosomes in the EquCab2.0 horse genome assembly on the left, and the boundaries of the 31 chromosomes in the assisted donkey genome assembly on the right. The horse chromosomes are ordered from #1 to #31, followed by chromosome X. The donkey chromosomes are ordered from largest (#1) to smallest (#30) autosome, followed by chromosome X. The dashed ovals shown in the two chromograms after the sex chromosome represent unanchored portion of the two genome assemblies. For ease of comparison, the orientation of donkey chromosomes was adjusted to match the orientation of the corresponding horse chromosomes. For "composite" chromosomes, the orientation of the horse chromosome that provided the largest amount of sequence was used when assigning orientation. An interactive version of the contact maps shown in the figure can be found at https://tinyurl.com/2hfwpruw.*

The output of the 3D-DNA/Juicebox Assembly Tools pipeline is a ".assembly" file that encodes the large-scale differences between the EquCab2.0 reference (2n=64) and the donkey (2n=62). The file is passed as input to Anonamos, a custom comparative assembly pipeline that examines per-base read alignment statistics to edit the reference at base-pair resolution at the same time incorporating larger rearrangements as annotated by the assembly file.

The Anonamos pipeline has the capacity to examine DNA-Seq and Hi-C alignment data (see Anonamos comparative assembler section). In order to simplify the comparison with the de novo donkey genome assembly (see *Domestic donkey de novo genome assembly section*) we used reads from the same donkey individual, Willy (*Equus asinus asinus*), for both the de novo and assisted genome assemblies. Specifically, the short-read DNA-Seq data from (Orlando et al. 2013) (SRA accession numbers SRR873443, SRR873444, SRR873445) aligned to EquCab2.0 (GCF_000002305.2) were used as input to Anonamos along with the ".assembly" file. The DNA-Seq reads were aligned using BWA-MEM (v 0.7.17-r1198-dirty, (Li and Durbin 2010; Li 2013)) and deduplicated using SAMtools (Danecek et al. 2021). The resulting EquCab2.0 sequence coverage was 15.6x. The alignment and deduplication statistics are included in Table S7.

**Anonamos comparative assembler.**

The Anonamos comparative assembler is a program that can generate a chromosome-length reference fasta from a set of reads, DNA-Seq and/or Hi-C, from an organism by mapping them to an assembly (draft or chromosome-length) of a closely related organism. Similarly to other tools of this type (see, e.g. AMOScmp (Pop et al. 2004) that has served as a primary inspiration for the tool, including its name) Anonamos attempts to substitute the traditional overlap-layout-consensus approach to assembly with alignment-layout-consensus. As part of the procedure, the read alignments to the assisting reference (provided in the form of .sam files, typically produced as part of the Juicer workflow) are examined in a base-by-base fashion to detect small-scale differences (single nucleotide polymorphisms, isolated small (compared to read length) deletions and insertions) between the two species. The identified differences are then incorporated into the assisting reference to create a consensus approximation of the organism represented by the read set.

Rearrangements between the two genomes pose the most difficult challenge to comparative assembly. For example, the analysis of read alignment signatures associated with putative rearrangements constitutes the most complex part of the AMOScmp pipeline (Pop et al. 2004). Unlike other comparative assembly tools, Anonamos does not attempt to extract the information necessary to resolve rearrangements from the alignment data. Instead, it relies on the established Hi-C data analysis by the 3D-DNA pipeline and takes the results as input in the form of a .assembly file.

The pipeline is available on GitHub (https://github.com/aidenlab/anonamos) and includes the following steps:

(0) In the preliminary step the work space is organized and the assisting reference is preprocessed to create a reference map file. A map file is a text file format in which, similar to mpileup, each line represents a single genomic position.

(1) During the next step (-S sort) the SAM file(s) are merge-sorted into a master sorted SAM file. The master SAM is then split into manageable chunks. The split size is adjusted dynamically to result in a predefined number of files determined by jobcount parameter (unprompted).

(3) In the mapping step (-S map) a map file is constructed from split alignment data to record bases observed at each genomic position. Indels and read breaks positions with respect to the reference are also recorded.

(4) The results from individual chunks are merged during the merge step (-S merge). Reference map data is also added during this step. (Including the assisting reference when generating the consensus yields credibility to singleton reads in low-coverage regions and helps distinguish them from contaminants and sequencing errors.) Optionally clip data is filtered in this step if necessary to remove read clips that originate from restriction sites recognized by the restriction enzyme when processing Hi-C read alignments.

(5) The next step (-S reduce) generates preliminary consensus from the map.

(6) The pipeline includes an optional step to reexamine indel and clip features that were not incorporated during preliminary data consolidation and reassemble the corresponding regions (-S reassemble).

(7) In the last step (-S finalize) the rearrangement file (provided via -a|--assembly flag) is incorporated and the final FASTA is constructed that reflects both the local consensus and the rearrangement data.

The Anonamos pipeline has a number of drawbacks, most obviously its dependence on the assembly of a closely related species. Even with a closely related species some highly divergent regions (arguably the most interesting regions!) might not be assembled due to issues with read alignment. Also, the current pipeline relies on majority voting when generating consensus which has limited ability to handle indels and does not address several important evolutionary events such as insertions in the target (segments of DNA present in the target genome that do not have a counterpart in the assisting reference genome) or segmental duplication events. (Note that the latter appear as coverage anomalies and the corresponding sequences typically end up in the unanchored portion of the assembly on account of them being identified as problematic by the 3D-DNA rearrangement detection workflow.)

It is also worth noting the following. The pipeline examines the differences between the assisting reference and the data in a somewhat bifocal fashion leading to potential issues in the mid-range. Specifically, Anonamos is well equipped to search for small polymorphisms (those measuring less than a read length). The smallest, single-nucleotide differences are the easiest type of inter-species polymorphisms to detect and resolve. At the same time, rearrangements are the easier to detect the larger they are using Hi-C, i.e. the most readily resolved rearrangement-type differences are at the opposite extreme of the size range spectrum. (Note that when speaking of the scale associated with translocation-type rearrangement the number is a function of the length of the fragment being translocated and the 1D distance between the original and the translocated position. Small sequences that are translocated further away along the chromosome may be easier to detect than larger sequences that remain close to their original position as the signal associated with the latter rearrangement will be obscured by all sequences in the 1D neighborhood "talking" to each other in 3D.) Previously, we showed that 3D-DNA can use Hi-C at ~10x coverage to reliably identify contact frequency differences between sequences separated by over ~10kb (Dudchenko et al. 2017). While additional coverage can be used to push the limit of detection to smaller scales one should expect that some rearrangements corresponding to the scales between ~100bp and ~10kb may remain unresolved.

**Domestic donkey *de novo* genome assembly.**

For de novo donkey genome assembly we combined a draft genome assembly ASM303372v1 (GCA_003033725.1) from (Renaud et al. 2018) with PBMC Hi-C data described in the *Domestic donkey Hi-C data* section. ASM303372v1 draft genome assembly was generated using Chicago HiRise (a technology that relies on HMW DNA) providing scaffolds of subchromosomal size (scaffold N50=15Mb). The two Hi-C datasets were aligned to the ASM303372v1 reference using Juicer (version 1.5.6, BWA 0.7.15-r1140, (Durand et al. 2016)) with '-s MboI' flag. The alignment statistics as generated by the Juicer pipeline can be found in Table S8. The resulting merged_nodups.txt file was passed as input to 3D-DNA (Dudchenko et al. 2017) which was run with default parameters to perform misjoin correction of the draft sequences as well as anchor them to chromosomes, order and orient them. The resulting candidate chromosome-length genome assembly was further examined using Juicebox Assembly Tools (Dudchenko et al. 2018). Hi-C data aligned to the ASM303372v1 draft and the final chromosome-length genome assembly (ASM303372v1_HiC) is shown in Figure S6. For convenience of comparison with the assisted assembly, the final chromosome-length scaffolds are ordered and oriented to match the order and orientation in the assisted donkey genome assembly shown in Figure S5.

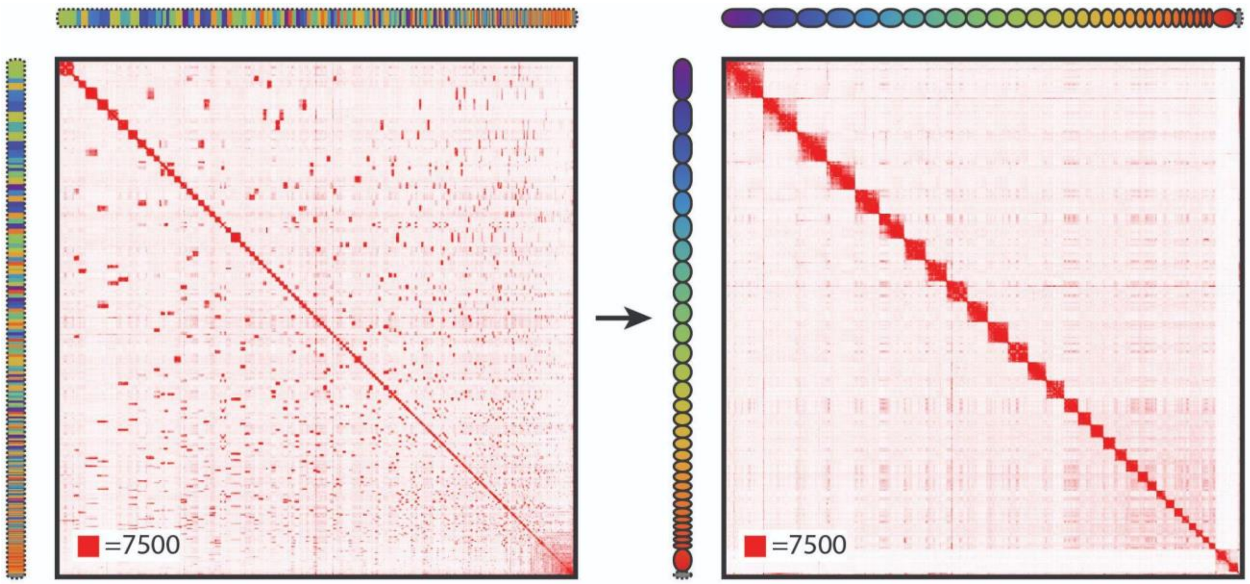


*Figure S6. Starting with the ASM303372v1 domestic donkey genome assembly draft from (Renaud et al. 2018), left, we used Hi-C data to error-correct, anchor, order and orient the draft sequences to produce a chromosome-length de novo assembly for the donkey, ASM303372v1_HiC, on the right. The contact maps show the same donkey Hi-C data, aligned to the draft on the left and lifted over to the chromosome-length reference on the right. Rainbow tracks on top and to the left of the contact maps are used to highlight corresponding loci between the two assemblies: the same color is used to show matching sequences. The draft sequences on the left are ordered by size, from largest to smallest. The chromosome-length sequences are ordered to match the ordering and orientation of the assisted donkey assembly (see* Domestic donkey assisted genome assembly *section). The ovals shaping the rainbow track on the right outline the boundaries of the 31 chromosomes in ASM303372v1_HiC. The dashed oval after the 31 chromosome-length scaffolds represents unanchored sequences. (The dashed oval around the draft assembly rainbow track highlights that all sequences in the draft are unanchored.) Interactive version of the contact maps shown in this figure can be found at https://tinyurl.com/25aaz94z.*

**Comparison of the assisted and de novo donkey genome assemblies.**

We compared the original horse, assisted donkey and de novo donkey assemblies by doing a whole-genome alignment using LASTZ (R. S. Harris 2007). The code was run with "--masking=3 --notransition --step=20 --nogapped --format=maf --ambiguous=iupac --hspthresh=50000" command options, and EquCab2.0 was used as a target and ASM303372v1 was used as query sequence. The resulting alignments were lifted over to ASM303372v1_HiC and to EquAsi_EquCab2.0_assisted_HiC to create dotplots shown in Figure S7.

Basic assembly statistics for the two donkey assemblies as well as the EquCab2.0 reference and ASM303372v1 can be found in Table S9. The stats were generated using BBMap (version 38.79) stats.sh script with default parameters (Bushnell 2014).

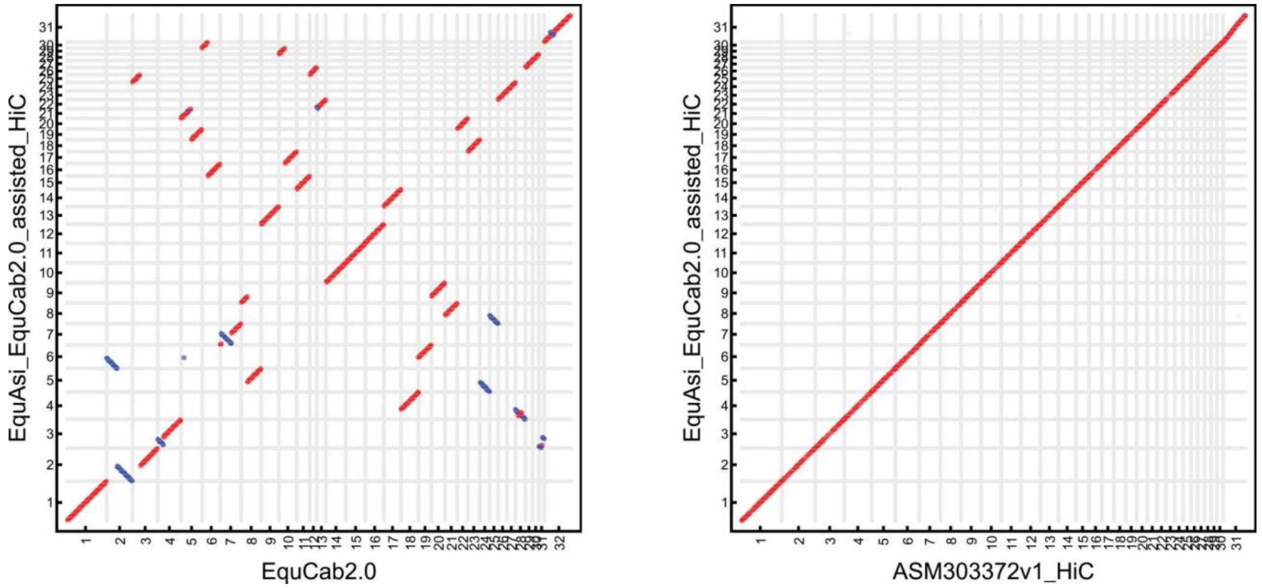


*Figure S7. Dotplots showing the correspondence between sequences in the chromosome-length assisted donkey genome assembly EquAsi_EquCab2.0_assisted_HiC and EquCab2.0, the assisting reference genome assembly (left); and a de novo chromosome-length assembly of the donkey, ASM303372v1_HiC (right). The left image was generated with liftover using the .assembly file describing a set of rearrangements between the donkey and the domestic horse identified from analyzing the donkey Hi-C data alignments to the horse reference. For the image on the right, the horse reference and de novo donkey sequences were aligned using LastZ alignment algorithm (R. S. Harris 2007) using "--masking=3 --notransition --step=20 --nogapped --format=maf --ambiguous=iupac --hspthresh=50000" command options, and the resulting alignments lifted over to the assisted donkey reference. A total of 15,000 highest-scoring alignment blocks are represented, with direct synteny blocks colored red, and inverted blocks colored blue. The chromosome order and orientation matches that in Figures S5 and S6. The chromosome borders are marked with vertical and horizontal gray lines. The dotplots illustrate excellent correspondence between the assisted and de novo assemblies despite many rearrangements occurring in the donkey and horse lineages after their split of the common* Equus *ancestor.*

#### Woolly mammoth assisted chromosome-length genome assembly.

The assisted pipeline was run as described in the previous section for the donkey. Specifically, the Hi-C data was aligned to the Loxafr3.0 reference using Juicer (version 1.6; BWA 0.7.17-r1188, (Durand et al. 2016)). The resulting deduplicated alignments were analyzed using 3D-DNA (Dudchenko et al. 2017) and Juicebox Assembly Tools (Dudchenko et al. 2018) to produce a ".assembly" file. The file was used as input to Anonamos, along with SAMtools deduplicated BWA-MEM alignments of aDNA-Seq data from the same individual published in (Díez-del-Molino et al. 2023). (The reads were subject to an adapter removal step prior to alignment.) The assisted assembly procedure resulted in 28 chromosome-length scaffolds (see Figure 2C). The autosomes were ordered from largest to smallest, and the sex chromosome identified as the chromosome having a reduced number of interchromosomal contacts, was placed last (as #28).

#### African elephant chromosome-length genome assembly.

To generate a chromosome-length genome assembly for an African elephant we used our previously published Hi-C dataset, SRA accession SRR19650826 (Álvarez-González et al. 2022). The data was generated from two fibroblast cell lines, one derived from a male, and another one from a female individual. For the purposes of this study the two datasets were processed separately. The data was aligned to Loxafr3.0 (Palkopoulou et al. 2018) and deduplicated using Juicer (Durand et al. 2016). The alignment and deduplication statistics are included in Table S10. Only female data was used for assembly, and both datasets were used for visualization purposes. The resulting merged_nodups.txt file was used as input to the 3D-DNA Hi-C guided assembly pipeline (Dudchenko et al. 2017) to produce a candidate genome assembly. The assembly was reviewed and polished using Juicebox Assembly Tools (Dudchenko et al. 2018). Contact maps generated with respect to the draft Loxafr3.0 genome assembly and the chromosome-length Loxafr3.0_HiC are shown in Figure S8.

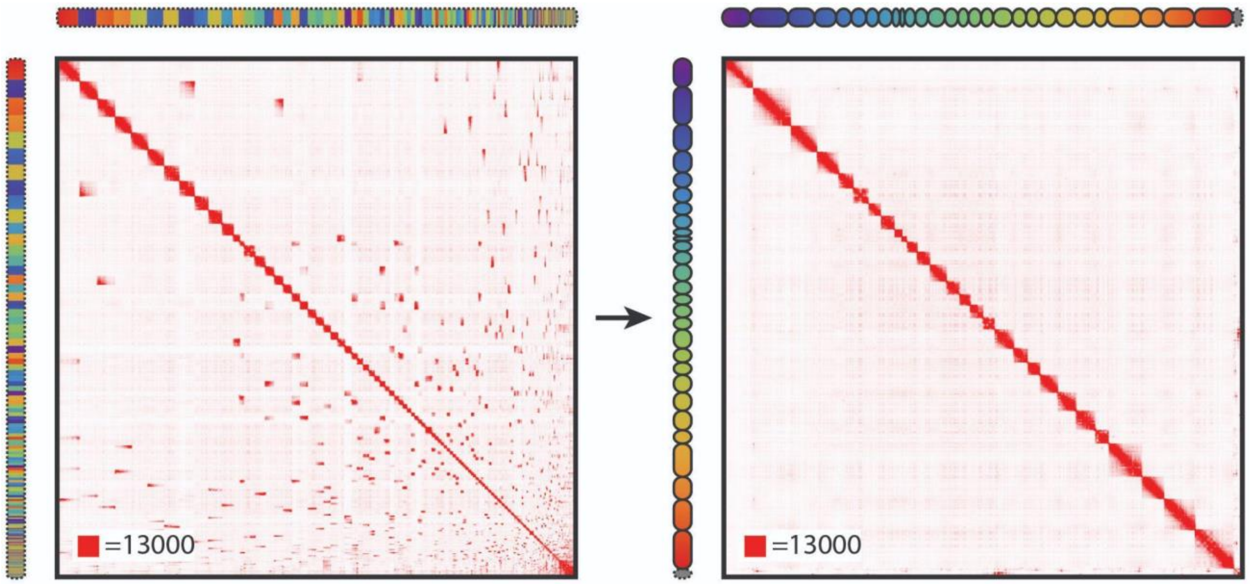


*Figure S8. Starting with a contact map for the African elephant draft genome assembly Loxafr3.0 from (Palkopoulou et al. 2018), left, we use in situ Hi-C data from (Álvarez-González et al. 2022) to error-correct, anchor, order and orient the draft sequences to produce a chromosome-length de novo assembly for the African elephant, Loxafr3.0_HiC, on the right. Rainbow tracks on top and to the left of the contact maps are used to highlight corresponding loci between the two assemblies: the same color is used to show matching sequences. The draft sequences on the left are ordered by size, from largest to smallest. The chromosome order on the right corresponds to the output generated by the 3D-DNA pipeline. The ovals shaping the rainbow track on the right outline the boundaries of the 28 chromosomes in Loxafr3.0_HiC. The dashed oval after the 28 chromosome-length scaffolds represents unanchored sequences. (The dashed oval around the draft assembly rainbow track highlights that all sequences in the draft are unanchored.) Interactive version of the contact maps shown in this figure can be found at* https://tinyurl.com/2c9ld7dv.

**Asian elephant chromosome-length genome assembly.**

We sequenced and used a combination of data from both female Asian elephant individuals in order to generate a chromosome-length genome assembly for the Asian elephant, a total of 5 libraries. The Hi-C sequencing data was aligned to ASM1433276v1 (GCA_014332765.1), a draft genome reference from (Tollis et al. 2021). The alignment was done using Juicer 1.6/BWA 0.7.17-r1188, run with the "-s none" flag (Durand et al. 2016). The resulting statistics for all libraries as generated by the Juicer pipeline are listed in Table S11.

The deduplicated alignments were used as input for the 3D-DNA pipeline (Dudchenko et al. 2017) and Juicebox Assembly Tools (Dudchenko et al. 2018) to perform misjoin correction, anchor, order and orient sequences in ASM1433276v1 and produce a chromosome-length reference ASM1433276v1_HiC. Contact maps built with respect to the original draft and the chromosome-length reference are shown in Figure S9.

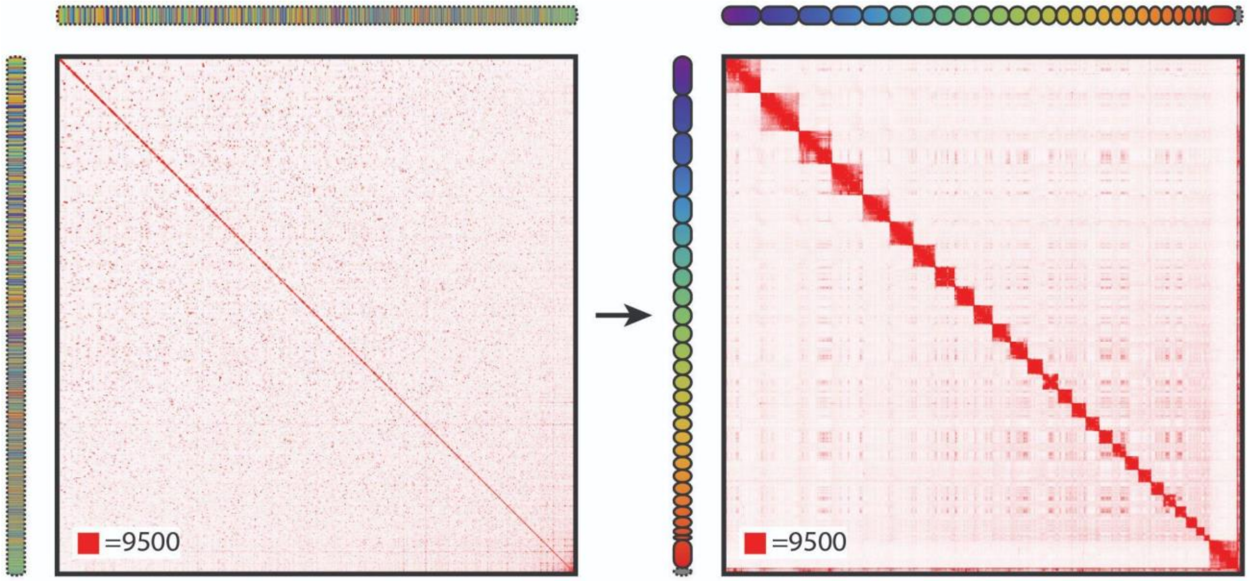


*Figure S9. Starting with the Asian elephant draft genome assembly ASM1433276v1 from (Tollis et al. 2021), left, we use in situ Hi-C data to error-correct, anchor, order and orient the draft sequences to produce a chromosome-length de novo assembly, ASM1433276v1_HiC, on the right. Rainbow tracks on top and to the left of the contact maps are used to highlight corresponding loci between the two assemblies: the same color is used to show matching sequences. The draft sequences on the left are ordered by size, from largest to smallest. The chromosome-length sequences on the left are ordered by side, from largest to smallest, except for the X chromosome identified via a characteristic interchromosomal contact pattern, which was placed last. The ovals shaping the rainbow track on the right outline the boundaries of the 28 chromosomes in ASM1433276v1_HiC. The dashed oval after the 28th chromosome-length scaffold represents unanchored sequences. (The dashed oval around the draft assembly rainbow track highlights that all sequences in the draft are unanchored.) Interactive version of this figure can be found at https://tinyurl.com/2c6euquk.*

### Synteny analysis in the Elephantidae family.

We compared the three Elephantidae genome assemblies by doing a whole-genome alignment using LASTZ (R. S. Harris 2007). The code was run with "--masking=3 --notransition --step=20 --nogapped --format=maf --ambiguous=iupac --hspthresh=50000" command options, and

Loxafr3.0 was used as target and ASM1433276v1 was used as query. The resulting alignments were lifted over to ASM1433276v1_HiC, Loxafr3.0_HiC and MamPri_Loxafr3.0_assisted_HiC to create dotplots shown in Figure S10. Chromosomes in ASM1433276v1_HiC, Loxafr3.0_HiC were ordered and oriented to match the order and orientation of sequences in MamPri_Loxafr3.0_assisted_HiC.

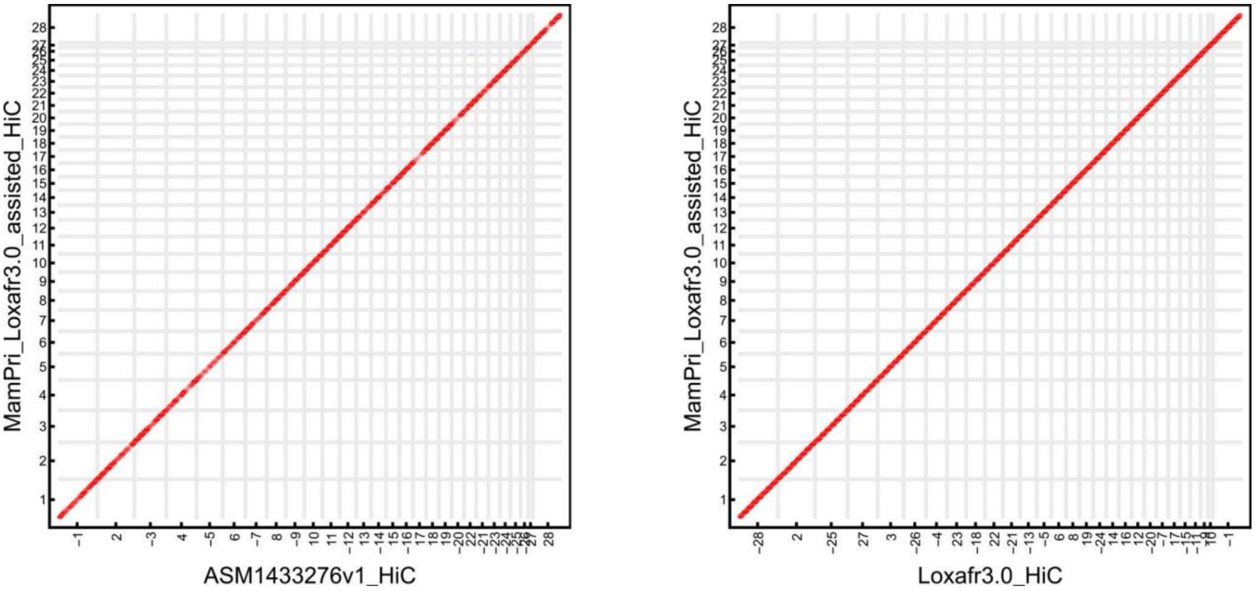


*Figure S10. Dotplots showing a conservation of synteny between the woolly mammoth and the Asian (left) and African (right) elephants. Each dot marks syntenic sequences between two genomes, with direct synteny blocks colored red, and inverted blocks colored blue. The syntenic blocks were identified using LASTZ alignment algorithm (R. S. Harris 2007). The pipeline was run with "--masking=3 --notransition --step=20 --nogapped --format=maf --ambiguous=iupac --hspthresh=50000" command options to align ASM1433276v1 to Loxaf3.0, and the resulting alignments lifted over to ASM1433276v1_HiC, Loxafr3.0_HiC and MamPri_Loxafr3.0_assisted_HiC. A total of 15,000 highest-scoring alignment blocks are represented. Chromosome borders are marked with vertical and horizontal gray lines. The chromosome order and orientation for ASM1433276v1_HiC and Loxafr3.0_HiC was adjusted to match that of MamPri_Loxafr3.0_assisted_HiC. Correspondence with original order and orientation is marked on the horizontal axis where each chromosomal block is marked with the original chromosome ID. A "-" sign before the ID indicates that the chromosome was inverted.*

**Woolly mammoth gene annotation.**

We used TOGA, a homology-based annotation pipeline that takes as input a gene annotation of a well-annotated reference species such as human and a whole-genome alignment between the reference and a query genome (Kirilenko et al. 2023). From the alignment data TOGA infers orthologous gene loci in the query genome, annotates and classifies them.

Prior to running TOGA, we masked repeats in the mammoth FASTA using MAVR (v0.1) (https://github.com/mahajrod/MAVR). MAVR is a wrapper to deploy three repeat annotation tools: RepeatMasker, version open-4.0.7 (Tempel 2012), Tandem Repeats Finder trf, version 4.04

(Benson 1999), and Windowmasker 1.0.0 (Morgulis et al. 2006). We then aligned the masked assisted mammoth assembly to the human reference hg38 (Schneider et al. 2017) using SegAlign (Goenka et al. 2020), a scalable GPU system for pairwise whole genome alignments based on LASTZ's seed-filter-extend paradigm. We customized SegAlign to take in the LASTZ "--inner" option (R. S. Harris 2007) and ran the pipeline with the following parameters: "--wga_chunk_size=2500000 --lastz_interval_size=4000000 --seq_block_size=40000000 --hspthresh=2400 --gappedthresh=3000 --ydrop=9400 --inner=2000". We then proceeded with creating alignment chains including identifying and incorporating repeat-overlapping alignments with (custom parallelized) RepeatFiller (Osipova, Hecker, and Hiller 2019) and removing chain-breaking alignments with chainCleaner (Suarez et al. 2017). The steps followed the recommendations outlined in https://github.com/hillerlab/GenomeAlignmentTools and https://github.com/ucscGenomeBrowser/kent/ (Kent et al. 2003).

The resulting chains were used as input to run TOGA (v1.1) with a "--mask_stops" flag using human annotations, U12 introns and isoform data shared at https://github.com/hillerlab/TOGA/tree/master/TOGAInput/human_hg38. We examined the resulting protein annotations using BUSCO v5.4.6 (Manni et al. 2021) which was run with a "-m proteins" flag and "-l eutheria_odb10" (Eutheria odb10.2021-02-19).

We ran the same annotation pipeline for the African elephant chromosome-length genome assembly Loxafr3.0_HiC. (The resulting annotations agreed well with the RefSeq gene annotations for Loxafr3.0 lifted to the chromosome-length genome assembly, data not shown.) The comparison of completeness of the resulting gene annotations, as a percentage of 11,366 eutherian genes from BUSCO, is included in Table S13.

**Compartment analysis.**

To simplify the comparison between the woolly mammoth skin and the Asian elephant data and to take advantage of the "golden standard" RefSeq annotations available for the African elephant (see *Genes associated with differential compartment calls*) all the compartment analysis was done with respect to the Loxafr3.0_HiC, the chromosome-length chromosome-length African elephant genome assembly. The maps were generated by lifting over alignments with respect to Loxafr3.0 (Table S1, S2). Chromosome IDs in Figure 3 are listed with respect to Loxafr3.0_HiC. Figure S10 can be used to lookup the syntenic chromosomes in the woolly mammoth and the Asian elephant genome assemblies.

The compartmentalization analysis largely follows (Lieberman-Aiden et al. 2009). The first order Pearson correlation analysis sharpens the signal, producing the characteristic plaid pattern that suggests that each chromosome can be decomposed into two sets of loci (traditionally labeled as A and B) such that contacts within each set are enriched, and contacts between sets are depleted.

The signal can be sharpened even further using second order Pearson autocorrelation analysis (Figure S11).

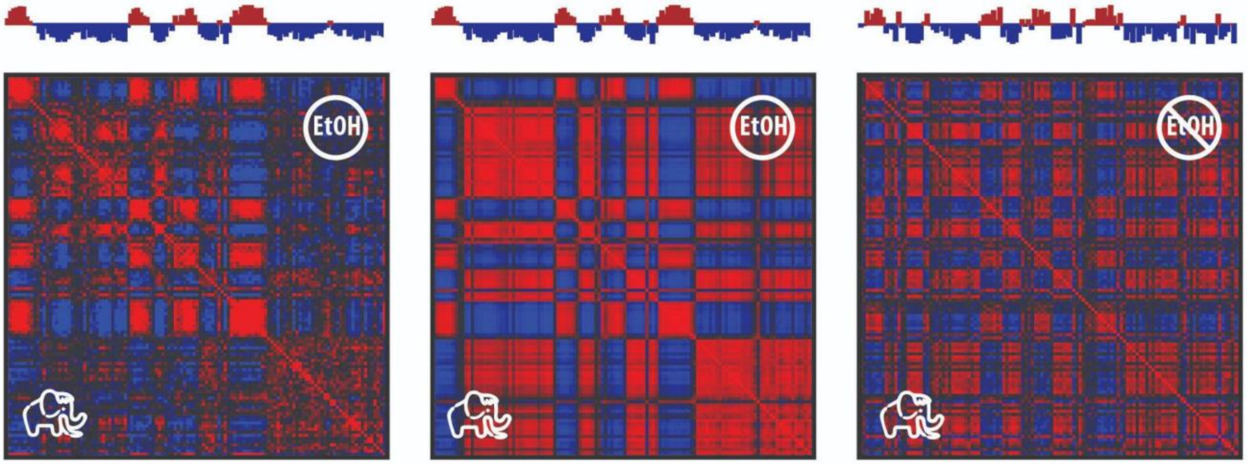


*Figure S11: PaleoHi-C contact maps from woolly mammoth skin exhibit a plaid pattern, characteristic of the spatial segregation between the A and B genome compartments. Here we show the 1st (left) and 2nd- (center) order autocorrelation matrices for the contact map from Figure 3A (chromosome 8 in mammoth syntenic to chromosome 23 in Loxafr3.0_HiC). The principal eigenvectors that can be used to distinguish between the A (active) and B (inactive) chromatin compartments are shown above the autocorrelation matrices. The matrices and eigenvectors are highly correlated, with the 2nd-order analysis resulting in sharper image due to stronger response to fine details in the map. The analysis can be performed even on suboptimal non-ethanol preserved sample data (right). While noisier, the 2nd-order Pearson map and activity profile match well the full dataset dominated by EtOH-preserved sample. All maps and tracks are shown at 1Mb resolution.*

The entries for the first autocorrelation matrix $p_{ij}$ were computed from observed/expected values ($c_{ij}$) for every intrachromosomal locus pair including i ($c_{xi}$) with every intrachromosomal locus pair including j ($c_{jx}$) except when x=i, x=j and i=j, and computing the Pearson correlation coefficient between the two resulting vectors. The KR-normalized observed/expected values for the purpose were extracted from the corresponding .hic files using Juicer Tools (Durand et al. 2016). For second order autocorrelation matrices, Pearson correlation coefficients between the two vectors representing all first-order Pearson correlation coefficients for loci i ($p_{ix}$) and j ($p_{jx}$) are calculated in the same fashion.

For the mammoth data, the first principal component of the KR-normalized contact matrix clearly corresponded to the plaid pattern for all but one chromosome (positive values defining one set, negative values the other) and was used to partition each chromosome into A and B. For HiC_scaffold_4, the second principal component corresponded to the plaid pattern. For elephant data the first principal component corresponded to the plaid pattern for all chromosomes.

We examined the distribution of values in the eigenvectors calculated for the mammoth and elephant skin contact matrices (Figure S12). While the elephant histogram reflected a typical bimodal distribution, the mammoth histogram did not show a binary A/B association reflecting, presumably, reduced confidence in assigning the relevant bins due to low data coverage in the mammoth. In view of this, we imposed a threshold of minimal absolute value of 0.05 for the assignment in the mammoth dataset to be considered reliable for downstream analysis.

As such, the cross-tissue comparison described in the text was done as follows. We first filtered out all bins for which the eigenvector value in the mammoth data was either positive and <0.05 or negative and > -0.05. We then applied the sign function to the remaining values and calculated corresponding Pearson coefficients. HiC_scaffold_4 was excluded from the analysis. If including the second principal component values for HiC_scaffold_4, the Pearson r values are as follows: elephant skin vs. mammoth skin: 0.920; vs. elephant ovary: 0.849; vs. elephant liver: 0.819, vs. elephant brain: 0.796; vs. elephant PBMCs: 0.758.

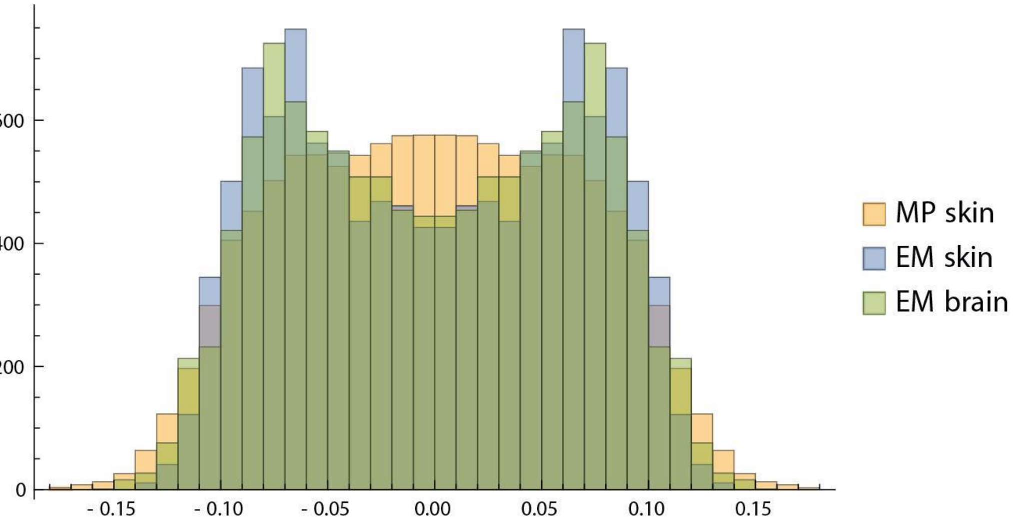


*Figure S12: A histogram of the eigenvector values for the mammoth skin (yellow), Asian elephant skin (blue) datasets and Asian elephant brain tissue (green). Because the eigenvector sign is arbitrary, both the eigenvector and values multiplied by -1 are included to build a symmetric plot. The elephant histograms show a typical bimodal distribution. The mammoth histogram does not display binary A/B association to the same extent. Instead, a large number of bins are assigned values close to 0. Reliable categorization presumably is not possible for the associated bins due to low data coverage.*

It is worth noting that the compartmentalization signal is robust enough that it can be observed solely in the suboptimal non-ethanol preserved supplementary dataset (Figure S11, right). The signal, while noisier, yields highly concordant segregation pattern and activity profiles for both the ethanol preserved and the suboptimal sample that was handled without preservation.

In addition to chromosome-specific Pearson-correlation maps shown in Figure 3 and S11 we calculated a genome-wide second-order Pearson correlation map for the woolly mammoth skin and the Asian elephant skin datasets. The matrix was calculated as described above, with observed/expected values for interchromosomal data extracted from the corresponding .hic files using Juicer Tools (Durand et al. 2016) along with intrachromosomal o/e, with genome-wide normalization. The resulting matrices demonstrate excellent agreement between the two datasets, for intra- and interchromosomal signals alike (see Figure S13).

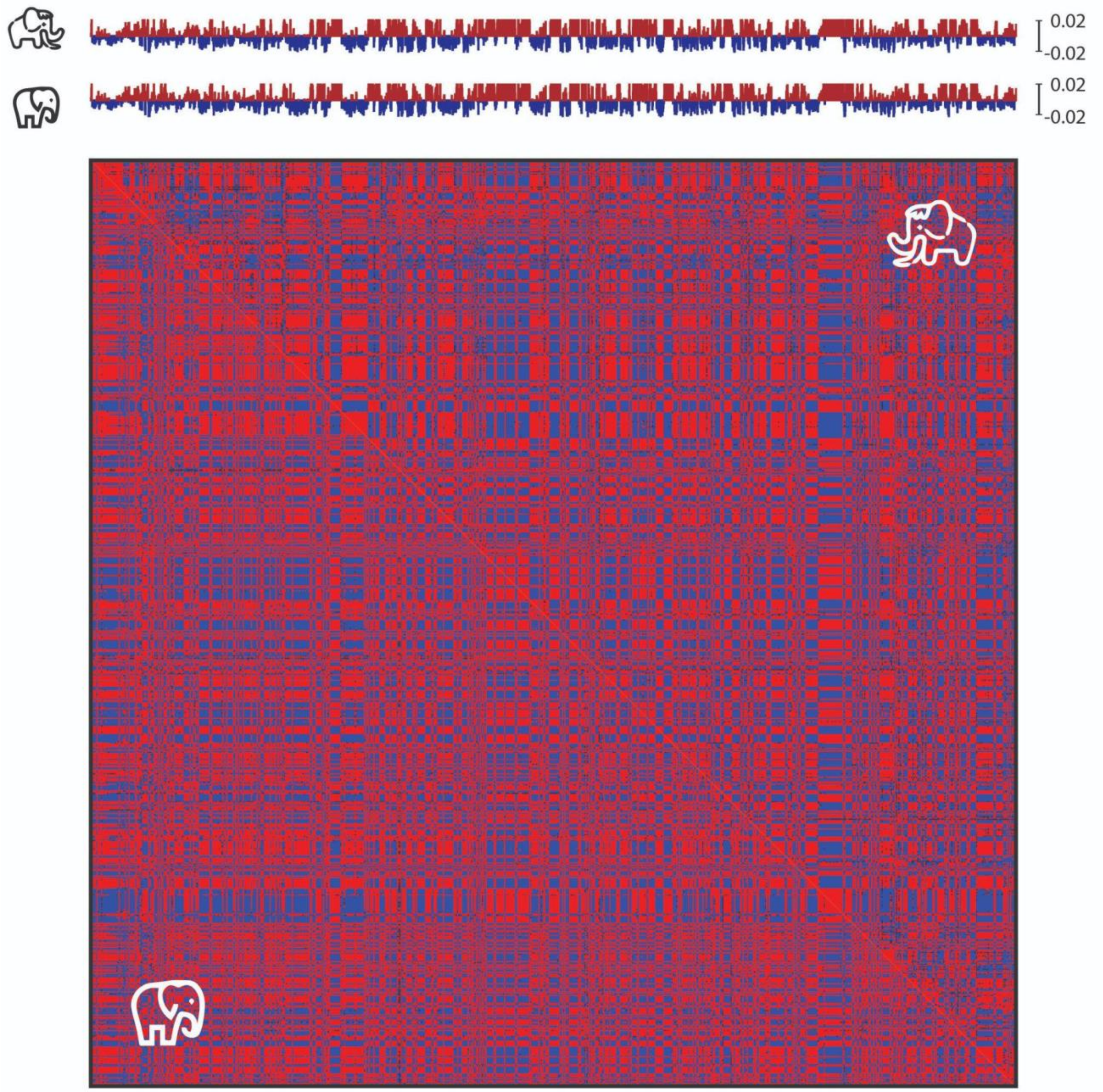


*Figure S13: Segregation into the active (A) and inactive (B) genome compartments is evident in the mammoth skin PaleoHi-C data genome-wide (upper right), and agrees extremely well with activity tracks computed for the modern Asian elephant skin contact dataset (lower left). The tracks above the Pearson maps show the first principal component of the elephant and the mammoth genome-wide contact matrices, with the woolly mammoth plotted on top, and the Asian elephant at the bottom. Pearson coefficient range shown is from -0.01 to -0.01 for both matrices. Eigenvector values range from -0.02 to 0.02 in both tracks.*

**Model and simulation details.**

We combined PaleoHi-C data with the OpenMiChroM (Oliveira Junior, Contessoto, et al. 2021) modeling package to construct an ensemble of 3D conformations for the woolly mammoth chromosome 8. The chromosome was represented using a polymer physics model, grounded in the maximum entropy principle. The model, known as the Minimal Chromatin Model (MiChroM), has been successfully used to study the organization of genomes in different organisms (Di Pierro et al. 2016; Oliveira Junior, Contessoto, et al. 2021; Oliveira Junior, Estrada, et al. 2021; Contessoto et al. 2023). The model accounts for differential interactions between the active and inactive (A and B) chromatin structural types.

The parameter values for the model were obtained from the mammoth Hi-C data aligned to Loxafr3.0_HiC binned at 250kb resolution, and chromatin types were assigned based on the first eigenvector from the experimental data autocorrelation matrix.

*Minimal Chromatin Model (MiChroM)*

The MiChroM energy function relies on two assumptions. The first is that the separation of chromosome phases is linked to the A/B (active and inactive) chromatin types. The second assumption is that the movement of different proteins is tied to how compact the polymer is. The MiChroM potential takes the following form:

$$U_{\text{MiChroM}}(\vec{r}) = U_{HP}(\vec{r}) + \sum_{\substack{k \geq l \\ k,\, l \,\in\, \text{Types}}} \alpha_{kl} \sum_{\substack{i \,\in\, \{\text{Loci of Type k}\} \\ j \,\in\, \{\text{Loci of Type l}\}}} f(r_{ij}) + \sum_{d=3}^{d_{cutoff}} \gamma(d) \sum_{i} f(r_{i,i+d})$$

where $U_{HP}(\vec{r})$ is the potential energy of a generic homopolymer; $\alpha_{kl}$ is the value of energy interactions between the chromatin type $k$ and $l$; $\gamma(d)$ is the ideal chromosome energy interaction as a function of the genomic distance $d$, and $f(r_{ij})$ represents the probability of crosslinking of loci $i$ and $j$. Similar to the approach for the human genome investigation (Di Pierro et al. 2016; Oliveira Junior, Contessoto, et al. 2021; Oliveira Junior, Estrada, et al. 2021), here it was also necessary to obtain the interaction energy parameters for each term of the potential $\alpha$ and $\gamma$.

The homopolymer potential employed here describes a generic bead-spring polymer in which each bead represents a genomic segment of 250 kb in sequence. The potential energy $U_{HP}(\vec{r})$ describes a spatially self-avoiding polymer and serves as a support for the features added by using the maximum entropy principle (Di Pierro et al. 2016).

This potential consists of the following four terms, $U_{\text{FENE}}$, $U_{\text{Angle}}$, $U_{hc}$ and, $U_{sc}$:

$$U_{HP}(\vec{r}) = \sum_{i\in\{\text{Loci}\}} U_{\text{FENE}}(\vec{r}_{i,i+1}) + \sum_{i\in\{\text{Loci}\}} U_{hc}(\vec{r}_{i,i+1}) + \sum_{i\in\{\text{Angles}\}} U_{\text{Angle}}(\theta_i) + \sum_{\substack{i,j\in\{\text{Loci}\}\\ j>i+2}} U_{sc}(\vec{r}_{i,j})$$

$U_{\text{FENE}}$ (Finite Extensible Nonlinear Elastic potential) is the bonding term applied between two consecutive monomers, connecting a sequence of beads with nonlinear springs. $K_b$ is the spring constant and $R_0$ is the equilibrium bond length.

$$U_{\text{FENE}}(\vec{r}_{i,j}) = \begin{cases} -\frac{1}{2}K_b R_0^2 \ln\left[1 - \left(\frac{r_{i,j}}{R_0}\right)^2\right] & r_{i,j} \leq R_0 \\ 0 & r_{i,j} > R_0 \end{cases}$$

Additionally, the hard-core repulsive potential $U_{hc}(\vec{r}_{i,j})$ is included to avoid overlap between bonded monomers:

$$U_{hc}(\vec{r}_{i,j}) = \begin{cases} 4\epsilon\left[\left(\frac{\sigma}{r_{i,j}}\right)^{12} - \left(\frac{\sigma}{r_{i,j}}\right)^6 + \frac{1}{4}\right] & r_{i,j} \leq \sigma 2^{\frac{1}{6}} \\ 0 & r_{i,j} > \sigma 2^{\frac{1}{6}} \end{cases}$$

A three-body interaction is included to all connected three consecutive monomers to consider a nonzero bending stiffness]. This angular potential that regulates the chain flexibility is given by:

$$U_{\text{Angle}}(\theta_i) = K_a[1 - \cos\theta_i - \theta_0]$$

where $\theta_i$ is the angle defined by two vectors $\vec{r}_{i,i+1}$ and $\vec{r}_{i,i-1}$.

All non-bonded pair interactions are described by a soft-core repulsive potential with the following form:

$$U_{sc}(\vec{r}_{i,j}) = \begin{cases} \frac{1}{2}E_{\text{cut}}\left[1 + \tanh\left(\frac{2U_{LJ}(\vec{r}_{i,j})}{E_{\text{cut}}} - 1\right)\right] & r_{i,j} \leq r_0 \\ U_{LJ}(\vec{r}_{i,j}) & r_0 \leq r_{i,j} \leq \sigma 2^{\frac{1}{6}} \\ 0 & r_{i,j} > \sigma 2^{\frac{1}{6}} \end{cases}$$

The expression $U_{LJ}$ corresponds to the Lennard-Jones potential:

$$U_{LJ}(\vec{r}_{i,j}) = 4\epsilon\left[\left(\frac{\sigma}{r_{i,j}}\right)^{12} - \left(\frac{\sigma}{r_{i,j}}\right)^{6} + \frac{1}{4}\right]$$

capped off at a finite distance, thus allowing for chain crossing at finite energetic cost (topoisomerases activity). $r_0$ is chosen as the distance at which $U_{LJ}(\vec{r}_{i,j}) = \frac{1}{2}E_{cut}$.

The model parameters are included in reduced units as:

$$K_a = 2\epsilon \quad K_b = \frac{30\epsilon}{\sigma^2} \quad E_{cut} = 4\epsilon \quad \epsilon = KT \quad R_0 = 1.5\sigma \quad \sigma = 1 \quad \theta_0 = \pi$$

*Crosslinking Probability Function*

One of the primary difficulties in developing chromosome models based on Hi-C maps lies in discerning the correlation between crosslink probability within the experiment and the geometric distance of a specific loci pair. MiChroM assumes that the likelihood of a loci pair $i$ and $j$ to form contact in 3D space decreases as a function of their distance $r_{i,j}$. In this sense, $f(r_{i,j})$ is adopted as a sigmoid function (switch function) designed to consider a high probability of crosslinking for short distances while long geometric distances should have a low probability of contact:

$$f(r_{i,j}) = \frac{1}{2}\left(1 + \tanh\left[\mu(r_c - r_{i,j})\right]\right)$$

where $\mu$ and $r_c$ values are determined based on the experimental Hi-C maps. The adjustment of the parameters considers two criteria: i) the function $f(r_{i,j})$ is calibrated to return 1 when two beads are in contact (distance between the center of two beads is equal to 1, in reduced units $\sigma$, i.e., $f(1) = 1$; ii) $f(r_{i,j})$ is also tuned to correctly return the experimental probability for the nearest neighbors where the maximum distance between the next nearest neighbors is 2 in reduced units $\sigma$. $f(r_{i,j})$ should decrease monotonically with the distance and the minimum value of the experimental probabilities must match with the following nearest neighbor maximum distance, i.e., $f(2) = \min\{P^{\text{exp}}_{i,i+2}\}$. The parameters adjusted for the Hi-C maps of Mammoth are $\mu = 2.27$ and $r_c = 1.7$.

*Energy Function Parameters from the Optimization*

The model parameters were obtained from the mammoth Hi-C data aligned to the Loxafr3.0_HiC genome assembly (HiC_scaffold_23 syntenic to mammoth chromosome #8). Data was binned at 250kb resolution. The parameters were optimized for the type-to-type interaction to accommodate for the A/B phase separation and for the ideal chromosome to consider the motor activity. A set of parameters that reflect the strength of A-to-A, B-to-B as well as A-to-B interactions is included in Table S14. The parameters are close to those estimated from human Hi-C data (Di Pierro et al.

2016). Some representative structures can be explored using the *Spacewalk* genome browser at https://tiny.3dg.io/PaleoHi-C-structures.

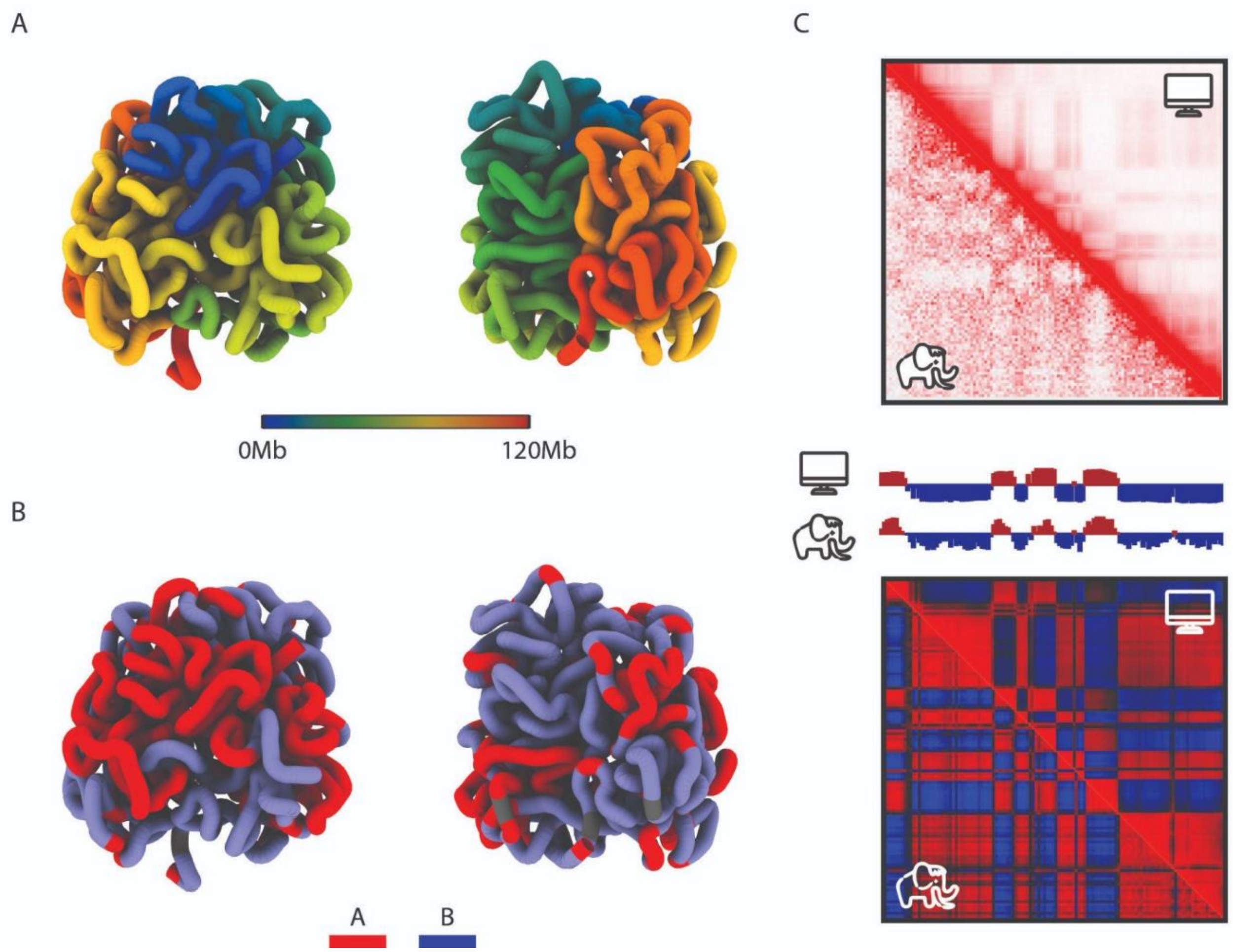


*Figure S14: Compartment assignments can be used to infer the structure of chromosomes in three-dimensional space by means of polymer physics simulations. A, B: Two representative structures from the simulated ensemble of 200,000 structures, colored by genomic index (A) and A/B types (B). C. Contact map generated from the ensemble of simulated 3D structures (top) and the corresponding Pearson's autocorrelation matrix (bottom). The simulated matrices (shown on the upper right) are compared with those calculated from PaleoHi-C data, shown in the lower left of the corresponding images. The simulated data shows a contact probability slope of -1.03 in the range between 1Mb and 10Mb genomic separation.*

**Genes associated with differential compartment calls in the mammoth and Asian elephant skin Hi-C datasets.**

We examined the A/B compartment annotations at 500Kb resolution for the mammoth and Asian elephant skin datasets to identify significant changes in compartmentalization between the two. Both the mammoth data and the Asian elephant data were aligned to the Loxafr3.0_HiC reference to take advantage of the RefSeq annotations available for the African elephant. The first principal component corresponded to the plaid pattern for all but two chromosomes at this resolution for both the mammoth and the elephant. These chromosomes (#4 and #10, the smallest chromosome) have been excluded from downstream analysis.

Similar to the analysis comparing the compartment assignments for different mammoth and elephant tissues, we ignored all positive bins of value <0.05 and negative bins of value > −0.05 (see Figure S12). We then proceeded with identifying 49 500Kb intervals that had differential compartment assignment for the mammoth and the modern Asian elephant skin dataset, i.e. either value ≥ 0.05 in mammoth and negative in the Asian elephant skin, or value ≤ −0.05 and positive in the Asian elephant. We used RefSeq gene annotations for Loxafr3.0 lifted to the chromosome-length genome assembly Loxafr3.0_HiC to look up the genes that reside in the differential intervals. The resulting gene set and the direction of change for the corresponding intervals are listed in Table S15.

**Supplementary compartment analysis using CRUSH (Compartment Refinement for the Ultraprecise Stratification of Hi-C).**

In addition to the eigenvector-based A/B categorization described above we used CRUSH, a tool for Compartment Refinement for the Ultraprecise Stratification of Hi-C, to do alternative type assignment analysis.

CRUSH is a modified version of the A/B index, a non-principal component analysis-based method for compartment identification published in (Rowley et al. 2017). In CRUSH, a subset of loci is initialized as pseudo "A" or "B", based on characteristics that correlate with and have been previously used to predict A and B compartments (Kalluchi et al. 2023; Zhou 2022). Because the A compartment contains high GC content and is gene-rich (Ramani, Shendure, and Duan 2016), using 500 bp bins, for each chromosome we define initialized pseudo-B as bins with GC content two standard deviations below the average. Pseudo-A are bins defined as bins overlapping genes and which do not have low GC content. Note that these initialized states represent only a small fraction of the chromosome, and that initial states are filtered and refined by actual interaction data from Hi-C experiments.

CRUSH then examines each row within the Hi-C matrix and measures the relative interaction intensities of that row with pseudo-A and pseudo-B similar to the A-B index (Rowley et al. 2017). This calculation includes distance normalization by (observed + 1) / (expected + 1), followed by z-scoring the signal for each row independently. Then the row's A-B index is calculated as the average z-score for interaction within pseudo-A minus the average z-score for interaction within pseudo-B. This initial calculation is performed to refine initialization states by removing pseudo-A bins that interact preferentially with pseudo-B, and vice-versa. Then the A-B index calculation is rerun with these new states and at multiple resolutions, refining pseudo-A and pseudo-B at a coarser resolution. For example, we refine initialized states at 1 Mb resolution and then use the refined states to initialize at 500 kb resolution. The resultant 1D track at each resolution can therefore be thought of as the row's relative interaction with features that correlate with A or B compartments.

We examined CRUSH output and compared the resulting A/B-type assignments with those from the eigenvector-based analysis at 1Mb and 500kb resolutions. The results are in excellent agreement (a Pearson correlation of ~0.9).

**Loop calls for aggregate peak analysis.**

We generated loop calls using HICCUPS (Juicer Tools version 1.9.9) (Rao et al. 2014; Durand et al. 2016) and published data from an African elephant fibroblast cell line (Álvarez-González et al. 2022) analyzed with respect to our chromosome-length African elephant genome assembly Loxafr3.0_HiC. The African elephant data was aligned to Loxafr3.0 (see Table S10) and lifted to Loxafr3.0_HiC. HICCUPS was run with default parameters using the contact map consisting of high mapping quality reads (mapq>=30) as input.

The resulting 3723 calls were further examined and rigorously filtered manually to remove 141 putative false positive calls to reduce a chance of false signal accumulation during aggregate peak analysis. Note that the loop count is lower than what is typically expected in a mammalian in situ Hi-C dataset (Rao et al. 2014) on account of the African elephant dataset being of relatively low depth (511,627,023 raw reads/345,636,788 Hi-C contacts, compare to 4.9 billion contacts used to generate loop calls for GM12878 human cell line in (Rao et al. 2014) which allows to identify only the strongest loops.

**Aggregate Peak Analysis.**

Juicer Tools (version 3.25.21) APA script was used to run aggregate peak analysis with KR normalization (-k "KR") (Durand et al. 2016). We ran APA for three Elephantidae datasets: African elephant fibroblast cell line data used to generate the loop calls, mammoth skin PaleoHi-C data and contact map generated from the modern Asian elephant skin sample, all using the same filtered loop list (see *Loop calls for aggregate peak analysis*). Note that the script filters the input loop list to exclude loops that are too close to the diagonal to be included at a given resolution.

All datasets were analyzed against the Loxafr3.0_HiC genome assembly to avoid the need to liftover loop positions, and only reads with mapping quality score mapq≥30 were used.

APA plots in Figure 4 of the main text are shown at 10kb resolution with 1883 of 3582 loops included in the calculation. Two additional resolutions (5kb and 25kb) are included in Figure S15.

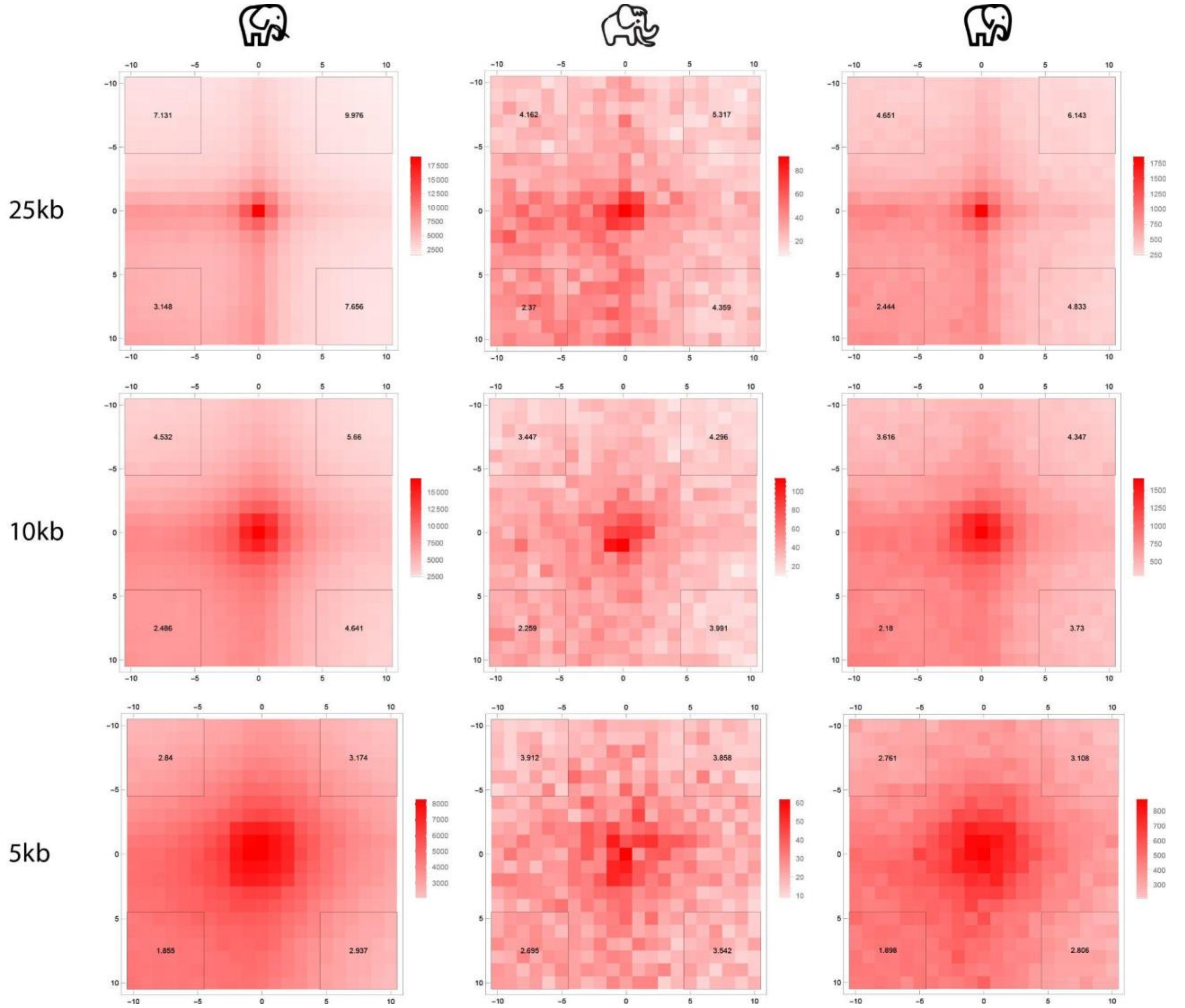


*Figure S15: APA plots for three Elephantidae datasets, the African elephant fibroblast data, mammoth skin PaleoHi-C and modern Asian elephant skin data, show enrichment of aggregate signal associated with the* Loxodonta africana *loop calls across multiple resolutions. Distance-filtered loop counts for the three resolutions are as follows: 492, 1880 and 3069 for 25kb, 10kb and 5kb, respectively.*

In order to further test against the possibility that the mammoth enrichment is brought about by an artifact we repeated the APA analysis using three random non-overlapping subsets of the African elephant loop calls, each 1000-call strong. The idea behind such analysis is that if, for example, the observed enrichment is associated with some a local “jackpot” effect where the number of contacts near one of the annotated loops is markedly increased in mammoth with respect to the background model due to an alignment artifact, we would expect that the enrichment in the central bin would only be observed in the subset that included the corresponding position and not the other two subsets. Instead, the analysis shows enrichment across all three subsets (APA score at 10kb resolution: 2.77, 2.65 and 1.92, respectively), consistent with the notion that the mammoth APA enrichments reflect real loop signatures.

**Estimating CTCF binding capacity along the X chromosome.**

We estimated CTCF-binding capacity along the X chromosome as follows. We extracted all PaleoHi-C reads where one end of the read aligned to the motif "CC.C[TC].G[AC]TGGCA.T". This is a consensus motif for the DXZ4/ICCE class of repeats that was manually curated based on a review of the literature (Horakova et al. 2012; Westervelt and Chadwick 2018). We then plotted the position of both the sequence matching read and its pair, on the logic that: a) if parts of the repeat sequence were assembled into the chromosome-length scaffolds the procedure would produce a hit in the correct position; and b) if the repeat sequence was not assembled or was not anchored, the second read would produce a hit to a nearby assembled portion of the chromosome-length scaffold. The procedure was designed to yield some robustness with respect to genome assembly quality, but can yield false positive hits, e.g. when a large number of repeats occur interspersed with a CTCF binding motif in one genomic position and without CTCF in another. The resulting chromosome-wide tracks for the X chromosome are shown in Figure 5. Peaks are visible at DXZ4, ICCE, and FROST. Peaks visible downstream of DXZ4 correspond to another CTCF-binding element, FIRRE (Darrow et al. 2016).

**ICCE boundary, just like DXZ4 boundary, disappears in males while FROST lies at the PAR boundary.**

We aimed to explore if, just like the DXZ4 boundary in humans, the boundaries identified on the mammoth X chromosome are specific to the inactive X chromosome. If this is indeed the case, we expect that the boundaries would be present only in female contact maps which show a combination of data from both the active, Xa, and inactive, Xi chromosomes. The boundary should disappear in contact maps generated from male samples, which only contain the Xa chromosome.

Figure S16 shows the male vs. female contact maps for the Asian and African elephants as well as in human. The images show disappearance of DXZ4 and ICCE boundaries in male elephants (Figure S16). The behavior of the boundary at FROST cannot yet be resolved in these datasets. As such, it is possible that FROST is not an Xi-specific element. See also the interactive version of the figure shared at https://tinyurl.com/22zgk8jn (note the inverted orientation of chrX in the African elephant chrX relative to the other chromosomes shown).

The analysis of male and female data near FROST can be used to show that the element flanks the PAR boundary. We observed that Hi-C coverage tracks for the X chromosome in both male and female Asian elephants are similar to one another from the p-terminus of the X chromosome until the position of this boundary element. On the opposite side of FROST, coverage is roughly 2-fold lower in males than in females (Figures S16). This is consistent with the hypothesis that the pseudoautosomal region (PAR) in mammoth and other elephantids extends to from the p terminus to FROST, and with earlier studies suggesting that the elephantid pseudoautosomal region is larger than its human counterpart (Raudsepp and Chowdhary 2015).

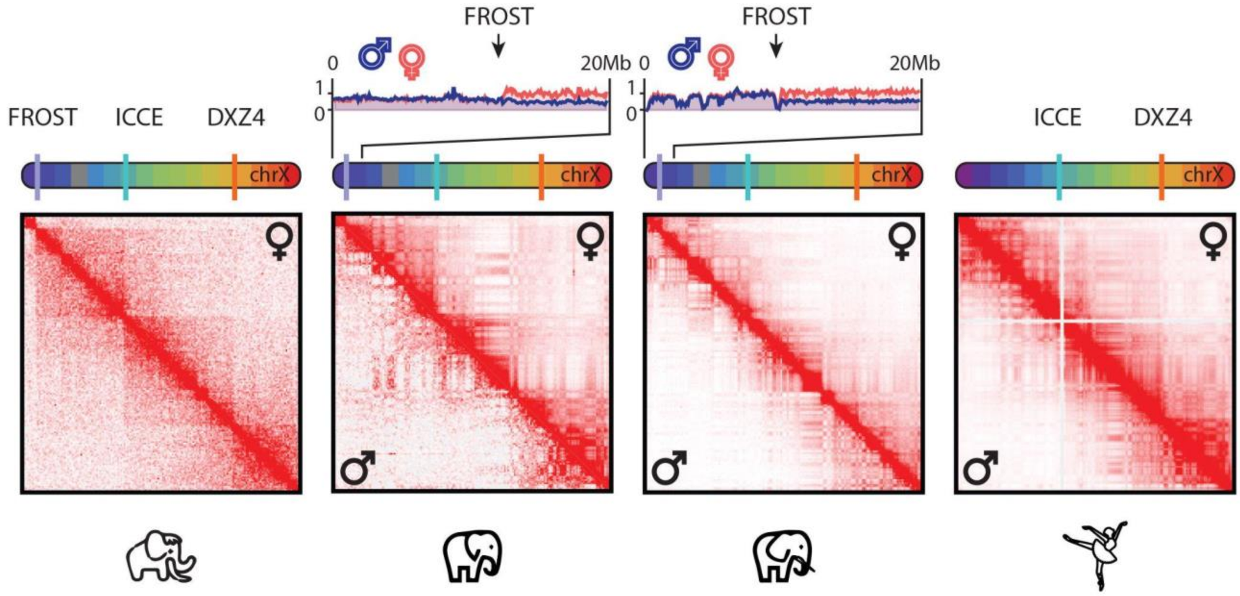


*Figure S16: ICCE boundary, just like DXZ4 boundary, disappears in males, while FROST flanks the pseudoautosomal region (PAR). The following chrX contact maps are shown: the woolly mammoth (PaleoHi-C skin data for a female individual), the Asian elephant (PBMC data for a male and female animal, in the lower left and the upper right part of the image, respectively), the African elephant (fibroblast data for a male and female animal (Álvarez-González et al. 2022), in the lower left and the upper right part of the image, respectively), and human (male AK1 from (H. L. Harris et al. 2023) in the lower left, and female GM12878 lymphoblastoid data from (Rao et al. 2014) on the right. Data is aligned to the following genome assemblies: MamPri_Loxafr3.0_assisted_HiC, ASM1433276v1_HiC, Loxafr3.0_HiC and hg19. The chromosomes have been oriented to match the conventional orientation in human. The chromograms above the contact maps indicate the ordering of the loci. The color scheme is based on the human chrX, purple (p-terminus) to red (q-terminus); corresponding loci across species are shown using the same color. Purple, cyan, and orange ticks on the chromograms indicate the positions of the FROST, ICCE and DXZ4 repeat elements, respectively. DXZ4 and ICCE elements disappear in male datasets. The Hi-C coverage tracks centered at FROST are shown at the top for male and female Asian and African elephant datasets. At FROST the coverage ratio for the male vs. female samples changes 2-fold, consistent with FROST lying at the PAR boundary. An interactive version of this figure can be found at [https://tinyurl.com/22zgk8jn](https://tinyurl.com/22zgk8jn) (note the inverted orientation for the African elephant chrX).*
